# Brain tissue transcriptomic analysis of SIV-infected macaques identifies several altered metabolic pathways linked to neuropathogenesis, and Poly (ADP-ribose) polymerases (PARPs) as potential therapeutic targets

**DOI:** 10.1101/2020.05.21.109140

**Authors:** Carla Mavian, Andrea S. Ramirez-Mata, James Jarad Dollar, David J. Nolan, Kevin White, Shannan N. Rich, Brittany Rife Magalis, Melanie Cash, Simone Marini, Mattia C. F. Prosperi, David Moraga Amador, Alberto Riva, Kenneth C. Williams, Marco Salemi

## Abstract

**Background:** Despite improvements in antiretroviral therapy, human immunodeficiency virus type 1 (HIV-1)-associated neurocognitive disorders (HAND) remain prevalent in subjects undergoing therapy. HAND significantly affects individuals’ quality of life, as well as adherence to therapy, and, despite the increasing understanding of neuropathogenesis, no definitive diagnostic or prognostic marker has been identified.

**Results:** We investigated transcriptomic profiles in frontal cortex tissues of Simian immunodeficiency virus (SIV)-infected Rhesus macaques sacrificed at different stages of infection. Gene expression was compared among SIV-infected animals (n=11), with or without CD8+ lymphocyte depletion, based on detectable (n=6) or non-detectable (n=5) presence of the virus in frontal cortex tissues. Significant enrichment in activation of monocyte and macrophage cellular pathways was found in animals with detectable brain infection, independently from CD8+ lymphocyte depletion. In addition, transcripts of four poly (ADP-ribose) polymerases (PARPs) were up-regulated in the frontal cortex, which was confirmed by real-time polymerase chain reaction.

**Conclusions:** Our results shed light on involvement of PARPs in SIV infection of the brain and their role in SIV-associated neurodegenerative processes. Inhibition of PARPs may provide an effective novel therapeutic target for HIV-related neuropathology.

## Introduction

The advent of combination antiretroviral therapy (cART) resulted in a 50% decline in rates of AIDS-related deaths, and a 40–50% decrease in the incidence of human immunodeficiency virus (HIV)-associated dementia (HAD) (Maschke *et al*, 2000). Yet an estimated 50% of infected patients exhibit HIV-1 central nervous system (CNS) infection (Zhao *et al*, 2009), with approximately 30% of in people living with HIV (PLWH) progressing to some form of HIV-associated neurocognitive disorder (HAND) (Heaton *et al*, 2010). Even in HIV-infected individuals on combined anti-retroviral therapy (cART), low-level viral replication persists in the central nervous system (CNS) (Spudich, 2016). Residual viremia as a result of incompletely suppressive cART (Massanella *et al*, 2013; Massanella *et al*, 2012) is associated with low-level immune activation driving chronic inflammation (Klatt *et al*, 2013; Massanella *et al*, 2016). It has been shown that both HIV and Simian immunodeficiency virus (SIV) can enter the CNS during early stages of infection (Resnick *et al*, 1988; Strickland *et al*, 2014), and there is compelling evidence that the brain is a putative reservoir for HIV (Marban *et al*, 2016; Wallet *et al*, 2019). Persistent CNS infection and inflammation may contribute to the development of HAND (Valcour *et al*, 2012), which remains a major cause of morbidity among HIV-infected individuals. As HAND-related cognitive decline is exacerbated by age-associated neurodegeneration, the prevalence of HAND is only expected to escalate with cART-increased life expectancy (Fogel *et al*, 2015). Moreover, if therapy is interrupted, viral rebound is going to occur (Andrade *et al*, 2020; Palmisano *et al*, 2007; Saez-Cirion *et al*, 2013), and because HIV is able to replicate in the CNS, brain specific viral variants are found at rebound after interruption of cART (Gianella *et al*, 2016).

While progress has been made in understanding the pathophysiology of HAND and neurological complications of HIV acquired immunodeficiency syndrome (neuroAIDS) under conditions of high viral load, the host’s inflammatory responses to low-level chronic systemic infection and how this exacerbates neuronal injury and dysfunction in the brain are incompletely understood. Infection of Rhesus macaques (*Mucaca mulatta*) with simian immunodeficiency virus (SIV) in the absence of therapy offers a well-established animal model for the study of the relationship of HIV infection and neuropathogenesis (Lamers *et al*, 2015; Mallard and Williams, 2018; Strickland *et al*, 2014), while avoiding the confounding factor of cART (Hatziioannou and Evans, 2012; Murray *et al*, 1992; Williams *et al*, 2008). Approximately 30% of Rhesus macaques infected with the heterogeneous SIVmac251 viral swarm (Strickland *et al*, 2011) develop within 2-3 years (Budka, 1991; Wiley *et al*, 1999) SIV-associated encephalitis (SIVE), the pathological hallmark of neuroAIDS, which is diagnosed *post mortem* by the presence of virus and abnormal histopathology features, such as inflammation of brain tissues and formation of multinucleated giant cells. When animals are depleted of CD8+ lymphocytes using an anti-CD8+ antibody before virus inoculation (Cartwright *et al*, 2016), the incidence is elevated to >85% in less than six months. Thus, CD8+ lymphocyte depletion provides a useful, rapid disease model with increased incidence of brain infection and neuropathology (Schmitz *et al*, 1999; Williams *et al*, 2005).

Myeloid cells accumulate in the meninges and choroid plexus during early infection, and in the perivascular space and SIVE lesions in infected macaques during late infection (Nowlin *et al*, 2015). In particular, SIVE lesions are composed of CD68+ CD163+ macrophages during early infection, as well as SIV-infected macrophages recruited terminally during simian AIDS (SAIDS) (Campbell *et al*, 2014; Nowlin *et al*, 2015). SIV-induced products of activated macrophages and astrocytes lead to CNS dysfunction and disease that might directly damage neurons (Roberts *et al*, 2003). These observations indicate that neuropathogenesis of HIV infection and pathogenesis of HAD and HAND may be linked (Kaul *et al*, 2005). It has also suggested that, given the neuroprotective properties of poly(ADP-ribose) polymerases (PARPs) inhibitors (Szabo *et al*, 2006), these inhibitors might be used as neuroprotective against NeuroAIDS as well (Rumbaugh *et al*, 2008). PARPs regulate a vast variety of cellular processes (Bai, 2015), and in particular, PARP1 and PARP-2 participate in regulating DNA metabolism (Ame *et al*, 2004), including DNA repair activated by DNA strand breaks (Morales *et al*, 2014). Previous studies demonstrated that PARP1 plays a role of in regulating HIV replication and integration (Ha *et al*, 2001; Kameoka *et al*, 2004).

Based on the hypothesis that PARPs appears to play an important role in HIV infection, we investigated the transcriptome of SIV-infected macaques with and without detectable virus in the brain to investigate whether PARPs expression is associated with SIV neuropathogenesis and biological processes translatable to HIV brain infection. We focused our analysis on characterizing the transcriptome profiles of the frontal cortex, as severity of cognitive impairment has been previously associated with the degree of frontal cortex neurodegeneration (Moore *et al*, 2006; Woods *et al*, 2009). In what follows, we report, for the first time, significant dysregulation of PARPs expression in SIV-infected brain tissues with detectable virus, associated with neurodegenerative processes.

## Methods

### Animal cohorts and sample collection

Frontal cortex tissue samples were collected from two cohorts of animals intravenously infected with SIVmac251 (Strickland *et al*, 2011), which originally consisted of five CD8+ lymphocyte-depleted (D) and six non-CD8-depleted (N = naturally progressing to SAIDS) Rhesus macaques, as previously described (Table 1) (Rife *et al*, 2016). Procedures on the CD8+ lymphocyte-depleted and naturally progressing cohort were conducted with the approval of New England Regional Primate Center at Harvard (Lamers *et al*, 2015) and University Tulane University’s Institutional Animal Care and Use Committee (Rife *et al*, 2016), respectively. Animals were kept in the same facility under similar conditions to minimize batch effects. Additional information on the treatment and handling of macaques in this cohort can be found in the study of Strickland et al. (Strickland *et al*, 2012). Gross pathology of the naturally progressing animals can be found in Rife et al. (Rife *et al*, 2016), and of the CD8+ lymphocyte-depleted ones in Table 1. All tissues collected during necropsy, following SAIDS onset and humane sacrifice, with the exception of animals N06, N07, D01, and D02, which were euthanized at 21 days post-infection (DPI) (Rife *et al*, 2016) (Table 1), were snap frozen in optimal cutting temperature medium and stored at −80° C. A single 50-100 mg section of frontal cortex tissue was used for RNA isolation. Viral DNA was extracted from frontal cortex tissues and detected by single genome sequencing (SGS) of the SIV envelope gene sequence as previously described (Rife *et al*, 2016; Strickland *et al*, 2014).

**Table 1.**
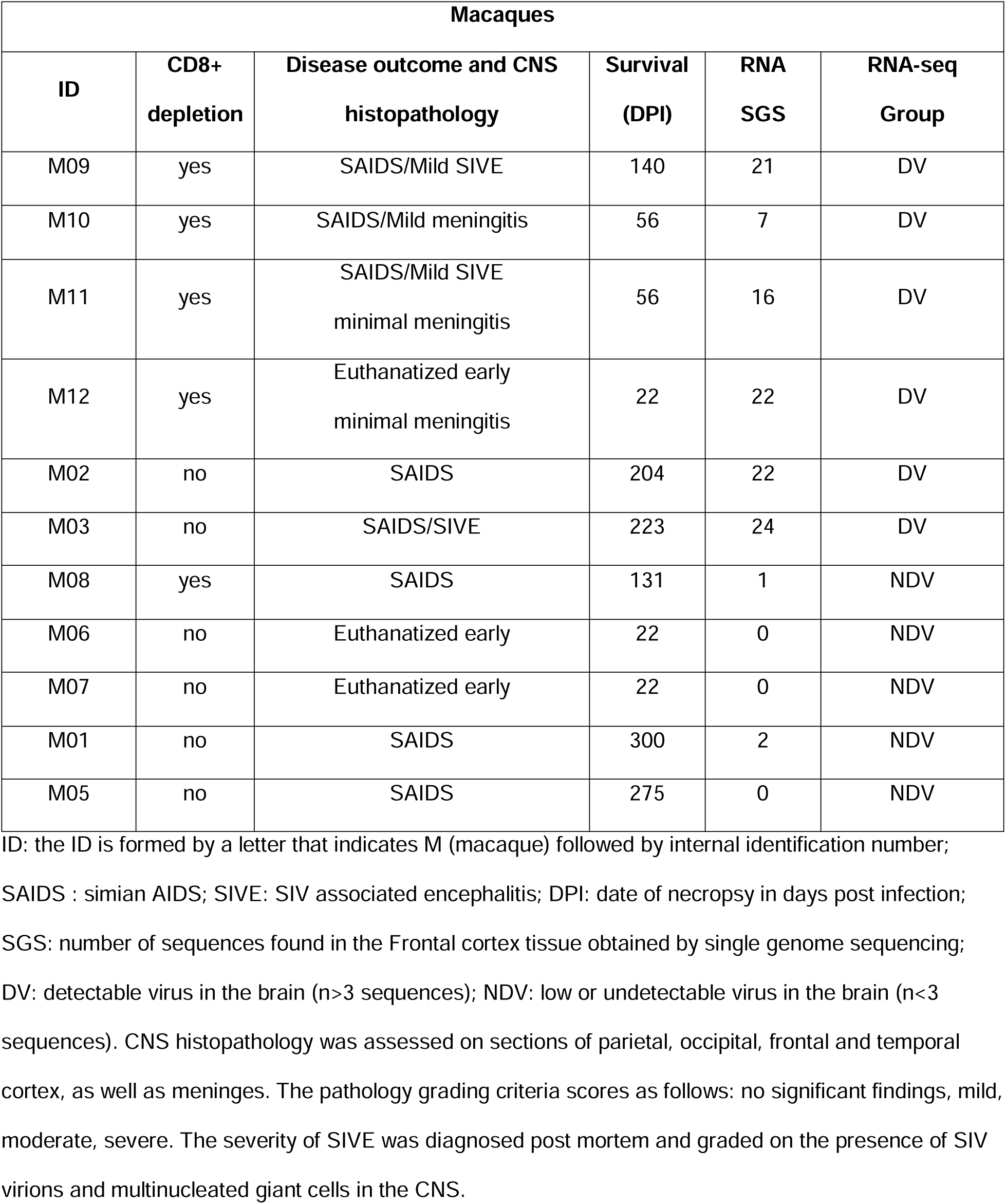
Epidemiological information on Macaques, Infection status by SGS.

| **Macaques** |  |  |  |  |  |
| --- | --- | --- | --- | --- | --- |
| **ID** | **CD8+**<br>**depletion** | **Disease outcome and CNS**<br>**histopathology** | **Survival**<br>**(DPI)** | **RNA**<br>**SGS** | **RNA-seq**<br>**Group** |
| M09 | yes | SAIDS/Mild SIVE | 140 | 21 | DV |
| M10 | yes | SAIDS/Mild meningitis | 56 | 7 | DV |
| M11 | yes | SAIDS/Mild SIVE<br>minimal meningitis | 56 | 16 | DV |
| M12 | yes | Euthanatized early<br>minimal meningitis | 22 | 22 | DV |
| M02 | no | SAIDS | 204 | 22 | DV |
| M03 | no | SAIDS/SIVE | 223 | 24 | DV |
| M08 | yes | SAIDS | 131 | 1 | NDV |
| M06 | no | Euthanatized early | 22 | 0 | NDV |
| M07 | no | Euthanatized early | 22 | 0 | NDV |
| M01 | no | SAIDS | 300 | 2 | NDV |
| M05 | no | SAIDS | 275 | 0 | NDV |

ID: the ID is formed by a letter that indicates M (macaque) followed by internal identification number;
SAIDS : simian AIDS; SIVE: SIV associated encephalitis; DPI: date of necropsy in days post infection;
SGS: number of sequences found in the Frontal cortex tissue obtained by single genome sequencing;
DV: detectable virus in the brain ( $n > 3$ sequences); NDV: low or undetectable virus in the brain ( $n < 3$
sequences). CNS histopathology was assessed on sections of parietal, occipital, frontal and temporal cortex, as well as meninges. The pathology grading criteria scores as follows: no significant findings, mild, moderate, severe. The severity of SIVE was diagnosed post mortem and graded on the presence of SIV virions and multinucleated giant cells in the CNS.

### RNA isolation and Next Generation Sequencing (RNA-Seq)

Total RNA was extracted with Qiagen RNeasy Lipid Tissue Mini Kit (Cat No: 74804) according to manufacturer protocol. Quantity and quality of RNA, from *post mortem* frontal cortex tissue samples, was assessed using the Invitrogen Qubit 2.0 and Agilent Tapestation 2200, respectively. Frontal cortex RNA sequencing libraries were prepared with Illumina TruSeq Stranded mRNA HT kit and sequenced on the 2×100 paired-end Illumina NextSeq platform at the University of Florida Interdisciplinary Center for Biotechnology Research.

### RNA-Seq data and pathway analysis

Paired-end reads were trimmed using trimmomatic (v 0.36) (Bolger *et al*, 2014), and quality control on the original and trimmed reads was performed using FastQC (v 0.11.4) (Brown *et al*, 2017). Trimmed paired-end reads were mapped to the *Macaca mulatta* genome available at Ensembl (http://dec2015.archive.ensembl.org/Macaca_mulatta/Info/Index). Sequences were aligned with STAR (v2.6.1) (Dobin *et al*, 2013). Reads were submitted to the Sequence Read Archive with the BioProject PRJNA624871. We obtained an average of ∼30.5 million reads for each sample, with an average of 55.9% of the reads mapped to the reference genome (Table S1), in line with typical percentage of transcriptome mapping (Conesa *et al*, 2016) (Table S1). Gene expression was quantified using RSEM (v1.2.31) (Li and Dewey, 2011). Differential expression analysis was performed using DESeq2 (Love *et al*, 2014), using a fold-change threshold of 0.05 and a significance level P < 0.01 (Table S2). For disease association enrichment and pathway analysis we opted for a cut off of (log_2_ (Log2) Fold-Change (FC)) of 1 of Log2(FC)-1 and P-value <= 0.05 to detect up- and down-regulated DEGs, respectively, as the FC represents genes that experienced 100% increase in expression (Table S3 and S4). These analyses were performed using the Ingenuity Pathway Analysis (IPA) software (Quiagen) after importing the list of 152 up-regulated (cut off Log2(FC)1 and P-value <= 0.05) and five down-regulated DEGs (cut off Log2(FC)-1 and P-value <= 0.05) (Table S2). The -log(p-value) of the pathway indicated the significance of overlap of the genes observed and the ones in the pathway and is calculated using the Fisher’s Exact Test (Fisher, 1934). Prediction of activation or de-activation of a certain pathway is based on the z-score, using a z-score threshold of 1.3. Calculation of the z-score of a pathway, which assess the match of observed and predicted up/downregulation patterns, is based on comparison between the direction of the genes observed compared to direction of those same genes in the active state of the pathway (Kramer *et al*, 2014) (Table S3 and S4).

### Quantitative PCR (qPCR)

cDNA from frontal cortex was generated with Invitrogen Superscript IV and random hexamers according to manufacturer’s protocols, using aliquots from RNA isolated for RNA sequencing. Comparative qPCR was conducted in triplicate for each sample using Applied Biosystems TaqMan Universal PCR Master Mix (ThermoFisher Catalog number: 4304437) and probes (0.25 µM) for PARP9, PARP12, PARP14, and Glyceraldehyde 3-phosphate dehydrogenase (GAPDH). Comparative qPCR was conducted with a 10-minute hold at 95°C, followed by 45 cycles of 95°C for 15 seconds and 60°C for 1 minute on the Applied Biosystems 7500 Fast Real-Time PCR System. Each sample’s mean C_T_ value for each qPCR reaction was normalized by subtracting the sample’s mean C_T_ for GAPDH to generate ΔC_T_. A standard deviation for the qPCR reaction was normalized with the standard deviation of GAPDH: S_ADJUSTED_ = (S_PROBE_ ^2^ + S_GAPDH_^2^)^1/2^. ΔΔC_T_ was calculated for each sample by subtracting its ΔC_T_ value from the mean ΔCT value of the samples without detectable virus in the brain. The fold difference in reference to the group of macaques without detectable virus in the brain was calculated with 2 ^−ΔΔCT^ and error bars were calculated with 2 ^−ΔΔCT ± SADJUSTED^. Statistical significance was tested using a Welch’s t-test, the hypothesis that two populations have equal means (Welch, 1947).

## Results

### Transcriptomic profiles are independent of CD8+ lymphocytes depletion

SGS of SIV *env* gp120 detected viral sequences in frontal cortex tissue of eight out of eleven animals: all five of the CD8+ lymphocyte-depleted and three of the non-depleted ones (Table 1). Three animals, two non-depleted (M06, M07) and one depleted (M12), were sacrificed early, while the others were sacrificed at SAIDS onset. As expected, while survival for depleted animals tended to be shorter, with an average of 81 days post infection (dpi), non-depleted animals’ survival averaged 174 dpi (Table 1). The number of positive PCRs in brain tissues at end point dilution varied between seven and 24 in most animals, except for two animals, M08 and M09, were only one and two SIV sequences, respectively, were detected, suggesting low level of brain infection as previously shown (Rife *et al*, 2016). Macaques with seven or more SIV sequences in the frontal cortex were all diagnosed with SIVE or meningitis at necropsy, with the exception of M02 (Table 1). The exception was not surprising, since we have shown in a previous study that an important co-factor linked to neuropathogenesis is viral compartmentalization in the brain, i.e. the presence of an adapted neurotropic sub-population, which was absent in M02 (Rife *et al*, 2016).

For all animals, RNA-Seq of frontal cortex samples resulted in high coverage (Table S1). Comparison of gene expression profiles in the frontal cortex between CD8+ lymphocyte-depleted and non-depleted macaques showed perturbance of only one gene – the nerve growth factor (NGF) gene, which resulted under-expressed in depleted animals with a fold change of Log2(FC)=3.3 – indicating that animals could be grouped, for further comparisons, independently of depletion status. Analysis of transcripts normalized expression among macaques corroborated that depletion category was not the dimension distinguishing the expression (Figure S1).

### Elevated antiviral gene response in macaques with detectable virus in the brain

The depleted versus non-depleted category analysis revealed that macaques with < 3 sequences were clustering with macaques with no detectable sequences in the brain (Figure 1, Table 1). Therefore, in order to minimize gene expression noise within the data due to inter-animal variability, macaques’ gene expression profiles were separated on the basis of a cut-off of n>3 SIV sequences detected by SGS in the brain (Figure 1, Table 1). Based on this cut-off, two non-overlapping groups could be defined: macaques with detect (DV) or low/undetectable (NDV) SIV in the brain. Differential expression analysis between DV and DNV macaque groups identified 102 up-regulated, and two down-regulated, differentially expressed genes (DEGs) in DV macaques with (Table S2). One of the two down-regulated DEGs (Table S2), NPAS4 (Log2(FC)-1.2) is a synaptic plasticity-promoting gene (Margineanu *et al*, 2018) crucial for synaptic connections in excitatory and inhibitory neurons and neural circuit plasticity (Ramamoorthi *et al*, 2011). Among the 102 up-regulated DEGs (Table S2), EPSTI1 (Log2(FC)2.8) plays a role in ensuring M1 versus M2 macrophage differentiation (Kim *et al*, 2018); SLFN13 (Log2(FC)2.3) restricts HIV replication (Yang *et al*, 2018). An important function of microglia is the presentation of foreign antigens to T lymphocytes (Schetters *et al*, 2017). The DV macaque group exhibited over-expression of the MAMU-A (Log2(FC)1.9) and MAMU-A3 (Log2(FC)1.7) genes, comprising the major histocompatibility complex class IA in Rhesus monkeys (Table S2). These genes are linked to disease progression during SIV infection (Zhang *et al*, 2002) (Table S2). Further corroboration of the presence of virus in the brain was given by up-regulation of components of antiviral interferon response, such the type I interferon (IFN)-stimulated genes (ISGs) ISG15 (Log2(FC)4.2) (Jeon *et al*, 2010) and ISG20 (Log2(FC)4.7) (Weiss *et al*, 2018), as well as of DDX60 (Log2(FC)3.9), a promotor of RIG1-like receptor-mediated signaling (Miyashita *et al*, 2011) (Table S2). Another iconic pathways hallmark of the innate immune responses are the role of pattern recognition of bacteria and viruses (z-score = 3.1) and activation of IRF by cytosolic pattern recognition receptors (z-score = 1.9) pathways, pathways that results in the activation of innate immune responses after recognition of pathogen-associated molecular patterns (PAMPs), such as lipopysaccharide or nucleic acids, by a variety of pattern-recognition receptors (PRRs) (Mogensen, 2009) (Table S3).

**Figure 1.**
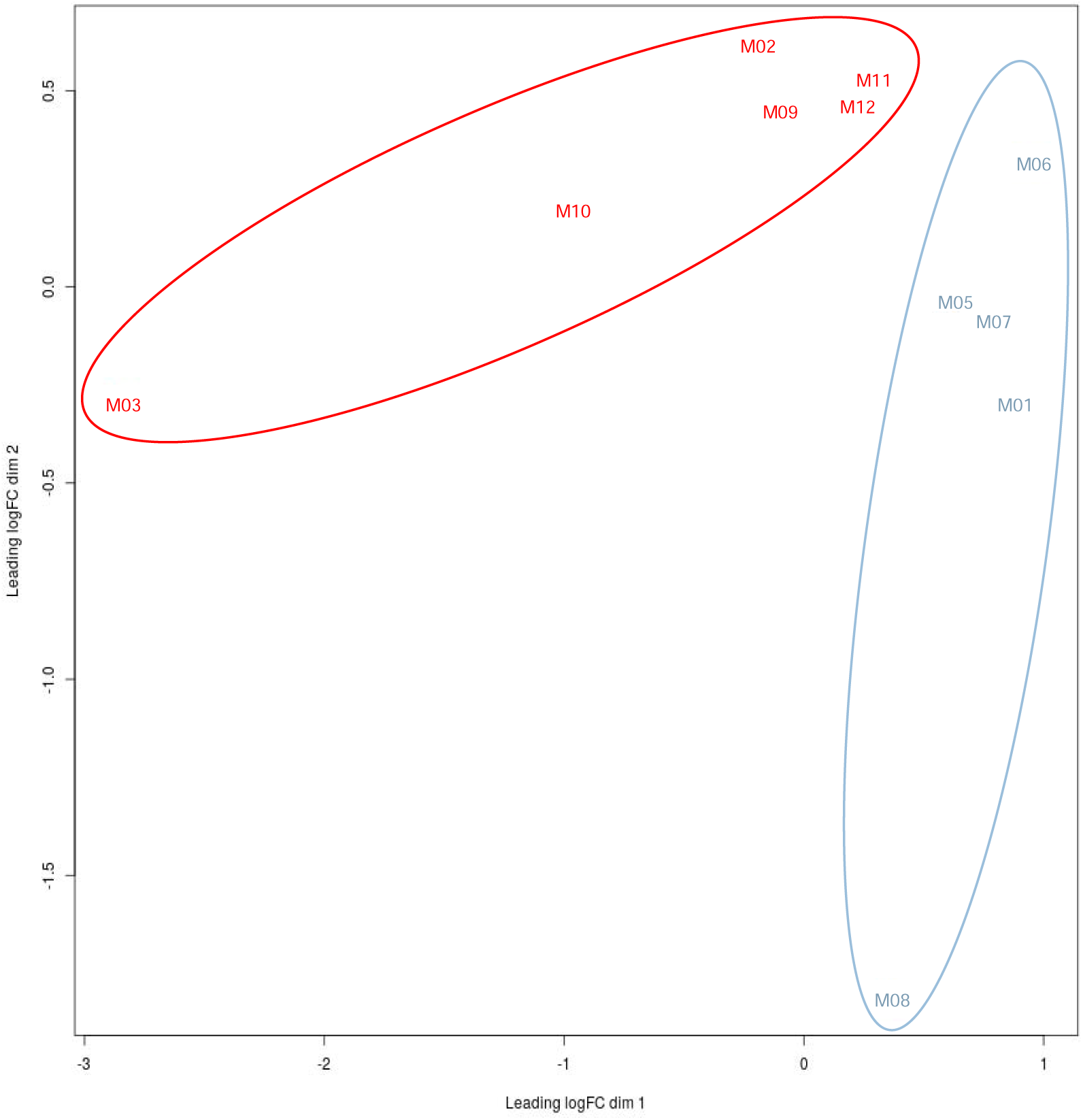
Multi-Dimensional Scaling (MDS) plot for the normalized expression data of DV and NDV SIV-infected macaques. Distance based the matrix of FPKM values quantified using RSEM v1.2.31 for all transcripts in all samples of macaques with detectable virus in the brain (n>3 sequences) in red, and macaques with low/undetectable virus in the brain (n<3 sequences) in blue. The plot shows good separation of the gene expression between the two groups, and non-overlapping.

### Orchestration of T cell apoptosis in brain through diverse pathways

Upon activation by T cell receptor and cytokine-mediated signaling, naive CD4+ T cells differentiate into types of T helper (Th) cells (Zhou *et al*, 2009), such Th1, playing a critical role in coordinating adaptive immune responses to various microorganisms interacting with CD8+ NK / CTL cells and macrophages (Romagnani, 1999). The inducible T-cell co-stimulator (iCOS) has been implicated in regulation of Th1, Th2 and Th17 immunity (Wikenheiser and Stumhofer, 2016) and plays an important role in recruiting entry of Th1 cells into inflamed peripheral tissue (Okamoto *et al*, 2004). In DV macaques several genes predicted the activation of the iCOS-iCOSL signaling in T helper cells pathway (z-score = 2), as well as of the Th1 pathway (z-score = 2.2) (Table S3). However, activation of T Cell Exhaustion Signaling Pathway (z-score = 1.3) was also predicted, which is characterized by loss of T-cell functions, that extendes to both CD8 and CD4 T cells (Yi *et al*, 2010) (Table S3). A lack of sufficient stimulation from secondary signals like cytokines - IL-12 and IFNγ are two important cytokines for Th1 differentiation that are not over expressed in our animals (Table S2) - may conversely lead to anergy or even apoptosis. Our animals exhibited activation of Calcium-induced T lymphocyte apoptosis pathway (z-score = 2), but also of Nuclear factor of activated T-cells (NFAT) (activation of NFAT in Regulation of the Immune Response pathway, z-score = 2.4) that is as an important mediator of T-Cell apoptosis (Table S3). These two pathways seems to be interrelated, as NFATs are calcium-dependent transcription factors, therefore activated by stimulation of receptors coupled to calcium-calcineurin signals (Park *et al*, 2020).

### Monocyte and macrophage activation in response to virus in the brain

Enrichment in activation of monocyte and macrophage cellular pathways (z-score = 2) was indicated by DEGs such as CD74 (Log2(FC)2.6), CD37 (Log2(FC)1.9), CSF1R (Log2(FC)1.2), and MNDA (Log2(FC)1.7) (Figure 2, Table S2 and S3). CSF1, in particular, has been associated with a positive feedback system wherein HIV infection increases CSF1 expression, followed by increased susceptibility of monocytes and macrophages to HIV replication upon exposure to CSF1 (Haine *et al*, 2006; Rappaport and Volsky, 2015). The fcy receptor-mediated phagocytosis in macrophages and monocytes pathway (z-score = 2) was also predicted to be activated (Figure 2, Table S2 and S3). Fc-mediated phagocytosis has been suggested as a successful mechanism for rapid control and clearance of HIV, as well as for reservoir eradication (Sips *et al*, 2016). Another pathway predicted to be activated lined to monocyte/macropahges activation is the TREM1 signaling pathway (z-score = 2.4) (Table S3). TREM1, a group of pattern recognition receptors, stimulates monocyte/macrophage-mediated inflammatory responses as its activation triggers expression and secretion of chemokines and cytokines that contribute to inflammation (Colonna and Facchetti, 2003). Additional evidence of activation of macrophages was given by the activation of the production of nitric oxide (NO) and reactive oxygen species (ROS) in macrophages pathway (z-score = 2.8), which allow for production of NO and ROS by activated macrophages, central to the control of infections (Forman and Torres, 2002) (Table S3).

**Figure 2.**
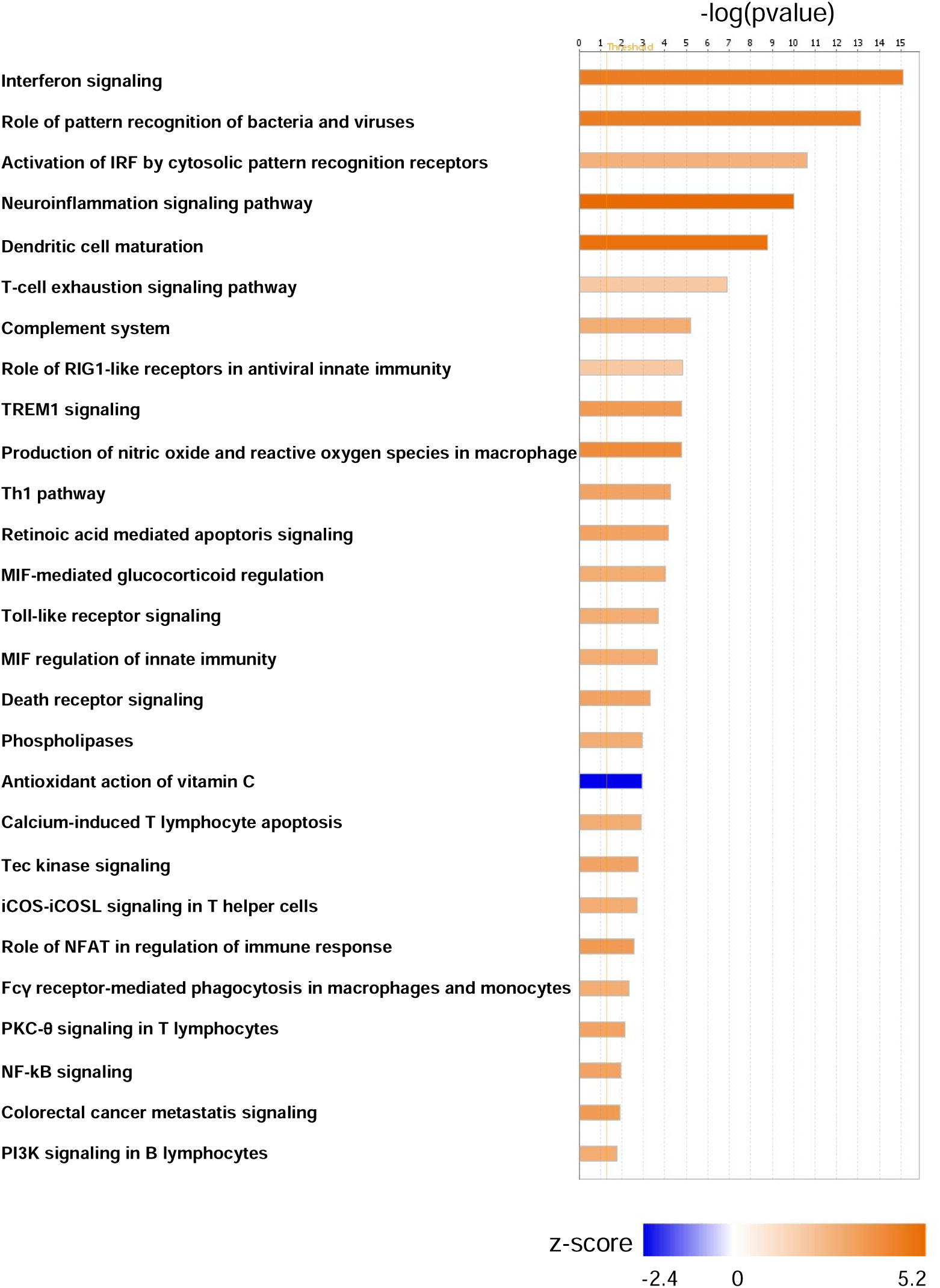
Predicted significant activated or de-activated intracellular molecular pathways from frontal cortex with SIV infection. Active or non-active state of pathways were predicted using the IPA library of canonical pathways, and significance was based on z-score greater than 1.3. The figure shows the number of genes being differentially expressed per pathway. In orange or blue is indicated the activation score (z-score).

Our results reflect previous transcriptomic studies that showed that the frontal cortex of SIV-infected macaques at terminal stage of SIVE was characterized by upregulation of STAT1, protein induced by cortical neurons, and ISG15, protein product of infiltrating macrophages (Roberts *et al*, 2003). Macrophage migration inhibitory factor (MIF)-regulation (z-score = 2) was also predicted to be activated (Table S3). MIF is a cytokine constitutively expressed by monocytes and macrophages in large amounts (Calandra and Roger, 2003) and an integral mediator of the innate immune system regulating host response through TLR4 (Roger *et al*, 2003), whereas TLRs initiate NF-κB and a number of other signaling pathways that broadly induce pro-inflammatory cytokines (Figure 2, Table S2 and S3) (Liu *et al*, 2017). Dysregulation of reactive oxygen species processes was indicated with NCF1 (Log2(FC)3.3), encoding for a NADPH oxidase that produces superoxide anions, inflammation, and organ injury through interaction with Toll-like receptors such as the DEG TLR4 (Log2(FC)1.49) (Gill *et al*, 2010) (Table S2 and S3).

### Inflammation as result of intensification of the innate immune response in presence of virus in the brain

Extending beyond the myeloid-mediated response, innate immunity pathways were identified as significantly differentiated, such Toll-like receptor (TLR) signaling pathways (z-score = 2) and interferon signaling (z-score = 3.2) (Table S3). The TLR signaling pathway was activated by up-regulation of CD14 (Log2(FC)1.64)), TLR3 (Log2(FC)1.94), and TLR4 (Log2(FC)1.49)) (Figure 2, Table S2 and S3). Genes that were upregulated in the interferon signaling pathways were: IFI35 (Log2(FC)1.0), IFI6 (Log2(FC)2.0), IFIT1 (Log2(FC)2.8), IFIT3 (Log2(FC)3.3), IRF9 (Log2(FC)1.8), ISG15 (Log2(FC)4.2), MX1 (Log2(FC)2.9), OAS1 (Log2(FC)2.5), PSMB8 (Log2(FC)3.3), STAT1, (Log2(FC)1.9) and STAT2 (Log2(FC)1.4) (Tables S2 and S3). In line to what previously reported during acute SIV infection in the brain of rhesus macaques, the interferon signaling pathway was predicated to be activated even in absence of high expression of either IFNα or IFNγ genes (Roberts *et al*, 2004) (Figure 2, Table S3). Intensification of innate immune response was also indicated by several DEGs, such C1QB (Log2(FC)2.0), C1QC (Log2(FC)2.3), and C3 (Log2(FC)1.5), involved in activation of complement and coagulation cascades (z-score = 2). Such complement cascades work to enhance the phagocytosis, proteolysis, inflammation, and overall magnitude of immune action (Janeway CA Jr, 2001). Complement system cascades have been linked to HIV-induced neurodegeneration in other research studies (Bruder *et al*, 2004; Speth *et al*, 2001) and to endothelial damage leading to reduced integrity of the blood brain barrier (Orsini *et al*, 2014) (Figure 2, Table S2 and S3). This increased innate immune response led to consequent up-regulation of numerous genes within the neuroinflammation signaling pathway (z-score = 3.6), likely establishing inflammation processes in the frontal cortex of the SIV-infected DV macaques (Table S3). Neuroinflammation signaling pathway plays a key role in maintaining the homeostasis of CNS, functioning to remove damaging agents, such SIV in this case, and clear injured neural tissues (Tohidpour *et al*, 2017). Excessive cell and tissue damage can ensue recruitment of microglia and enhancement of their activities, which exacerbates neuronal damage and ultimately results in chronic inflammation with necrosis of glial cells and neurons (Wang *et al*, 2015). Necroptosis is a regulated necrotic cell death pathway that defends against pathogen-mediated infections, morphologically characterized by the loss of cell plasma membrane and the swelling of organelles, particularly mitochondria. Compared to apoptosis, necroptosis generates more inflammation. Several death receptors promote necroptosis when activated, including tumor necrosis factor receptor TNFR1, Fas, TNFRSF10A and TNFRSF10B – with up-regulation of its ligand TNFSF10 (Log2(FC)1.13) – as well as TLRs (Feoktistova and Leverkus, 2015; Najafov *et al*, 2019) (Table S2 and S3). Activation of pathways associated with interferon (z-score = 3.2) and death receptor signaling (z-score = 2.2) are likely to be associated with neuronal apoptosis, similarly to what reported for infection of neurotropic West Nile virus in the brain (Clarke *et al*, 2014) (Table S3). Finally, neuronal damage was also suggested by up-regulation of PSMB8 (Log2(FC)3.3) and PSMB9 (Log2(FC)3.0), crucial for proteasome activity and regulation of protein turnover in neuronal synapses (Speese *et al*, 2003). PSMB8 and PSMB9 have been previously implicated in research studying SIVE-induced neuronal dysfunction (Gersten *et al*, 2009b) (Table S3). Lastly, NCF1 produces superoxide anions causing increased oxidative stress, which is linked to nervous system damage (Starkov *et al*, 2004; Uzasci *et al*, 2013), and activation of STAT1 (Log2(FC)1.9) provides further evidence of response to oxidative stress (Olagnier *et al*, 2014) (Table S3).

### Upregulation of PARPs in the frontal cortex of macaques with detectable SIV in the brain

Transcripts of four PARPs were up-regulated in the SIV-infected frontal cortex: PARP9 (Log2(FC)1.8), PARP10 (Log2(FC)1.9), PARP12 (Log2(FC)1.9), and PARP14 (Log2(FC)2.7) (Figure 3a, Table S2 and S3). Over expression of these PARPs was also corroborated by quantitative PCR (Figure 3b). Expression of PARP1, a member of the PARPs family that has been the focus of HIV research due to their role in viral integration, replication, and transcription (Bueno *et al*, 2013; Ha *et al*, 2001; Ha and Snyder, 1999; Hassa and Hottiger, 1999; Kameoka *et al*, 2004; Kameoka *et al*, 2005; Rom *et al*, 2015), as well the other PARPs, was not significantly over or under regulated (Table S5), as also confirmed by qPCR of mRNA transcripts (Figure 3b). PARPs are known to be activated by DNA strand breaks (Ikejima *et al*, 1990; Ray Chaudhuri and Nussenzweig, 2017), such ones occurring in HIV integration, as well as by interferon response (Atasheva *et al*, 2014). While there are mixed reports as to whether (Ha *et al*, 2001; Kameoka *et al*, 2005) or not such genes are necessary for HIV integration (Ariumi *et al*, 2005; Baekelandt *et al*, 2000), their function as a transcriptional repressor of HIV and inhibitor of cellular translation is known (Atasheva *et al*, 2014; Bueno *et al*, 2013). Upregulation of PARP9, PARP10, PARP12 and PARP14 and TNFSF10 predicts the activation of the death receptor signaling pathway (z-score = 2.2), associated with programmed cell death, and the retinoic acid mediated apoptosis signaling pathway (z-score = 2.2) (Figure 2, Table S2 and S3), which functions as an important regulatory signaling molecule for cell growth, differentiation and neurodegeneration (Das *et al*, 2014).

**Figure 3.**
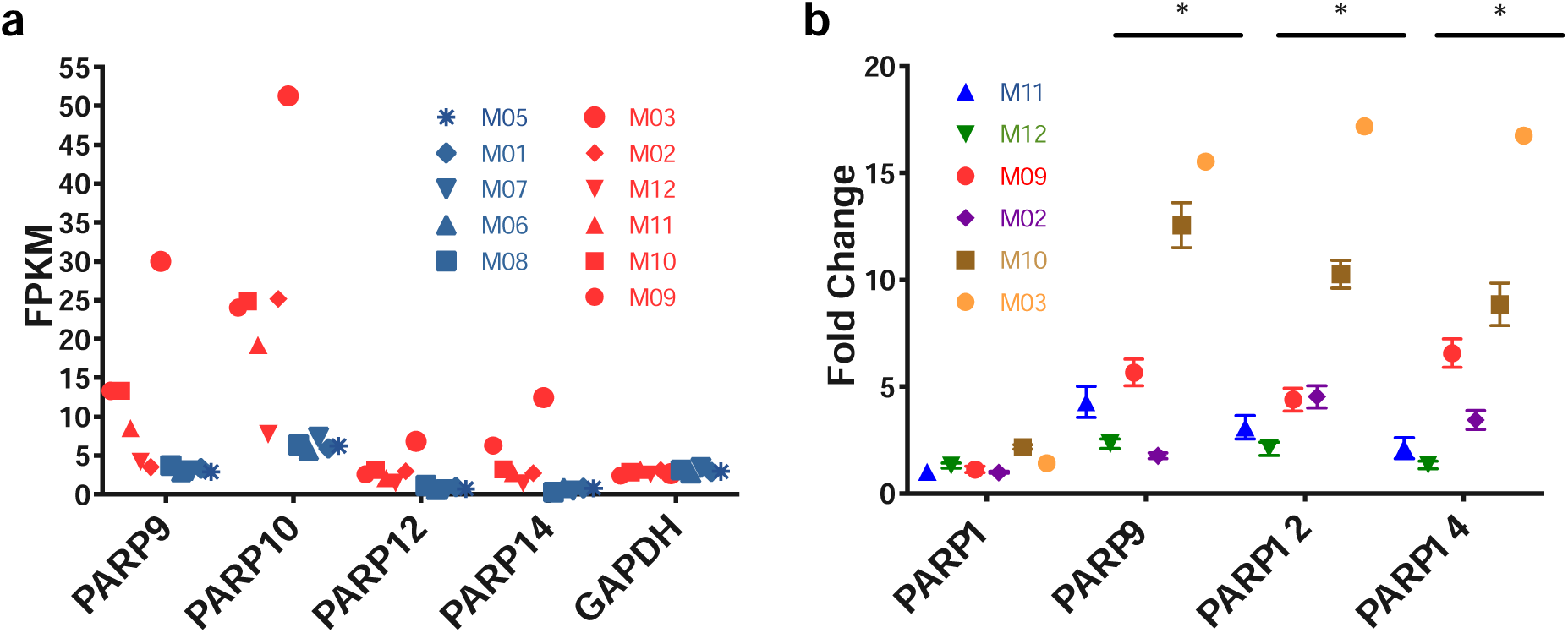
Differential expression of PARPs in the frontal cortex of macaques with detectable virus in the brain. (a) FPKMs (Fragments Per Kilobase of transcript per Million mapped reads) of PARP1, PARP9, PARP12 and PARP14 transcripts in macaques with detectable virus in the brain (red) and in macaques without detectable virus (blue). (b) Quantitative PCR analysis of mRNAs levels of PARP1, PARP9, PARP12 and PARP14 expressed in frontal cortex for macaques with detectable virus as relative to the averaged mRNA expression of the PARPs found in macaques without detectable virus. Colors indicate different macaques, while symbols are indicating the same macaque as shown in panel a. Asterisks indicate p < 0.001.

## Discussion

The CNS has gained importance as a potential reservoir during persistent HIV infections and the renewed focus of intense efforts on eradication strategies (Hellmuth *et al*, 2015; Salemi and Rife, 2016; Saylor *et al*, 2016). We have presented evidence of activation of pathways that implicate a significant myeloid response to SIV infection in the brain of a well-established model of HIV disease progression, even in macaques euthanatized early. We recognize that the present study has some limitations, as it lacks of un-infected animals as controls, and contrasted groups are mixed, including both naturally progressing and CD8+ depleted animals. As our goal is to understand how presence of the virus in the brain plays versus its absence during infection, the first limitation is easily overcome as by comparing infected and un-infected animals would not address our question. As for the second limitation, although it might seem counterintuitive that CD8+ depletion has no effect on the transcriptomics profiles of frontal cortex, it is important to remind that CD8+ depletion impact the peripheral circulation of CD8+ lymphocytes but not in meninges (Ratai *et al*, 2011), and that depletion alone does not have measurable effects on neuronal integrity preserving brain metabolism (Ratai *et al*, 2011). It is also noticeable that previous studied demonstrated that CD8-deplitoin does not alter metabolite levels, does not cause astrogliosis or microglial activation as compared to SIV-infected animals (Ratai *et al*, 2011). This last finding validates that neuroinflammation in these macaques is not dependent on depletion, but rather on presence of the virus in the brain. These findings confirm the validity of our approach, as that the predicted activated neurodegenerative pathways observed in our study are potentially due to the presence of virus and its manipulation of the immune system, rather than by absence of CD8+ T cells. The results agree with HIV and SIV entry in the CNS during early infection (Resnick *et al*, 1988; Strickland *et al*, 2014). Presence of virus in frontal cortex was linked to upregulation of gene expression, as well as neuropathology with the exception of animal M09 (Rife *et al*, 2016). It is interesting to note, however, that virus compartmentalization (distinct neurotropic subpopulation) in the brain, which has been linked to neuropathogenesis (Lamers *et al*, 2015; Mallard and Williams, 2018; Strickland *et al*, 2014), was also absent in this animal (Rife *et al*, 2016). Therefore, while virus induced dysregulation of gene expression seems to play an important role, the emergence of an SIV neurotropic sub-population may be a necessary condition for the onset of neuroAIDS, at least in the macaque model.

Akin to previous studies (Roberts *et al*, 2004), our findings indicate that frontal cortex of macaques with detectable SIV in the brain have significant upregulation of several genes. In particular, our results support that SIV in the frontal cortex alters transcriptional pathways associated with innate immune response, neuroinflammation, oxidative stress, and cellular death, interferon/STAT1 pathway, and monocyte/macrophage migration in line with previous studies (Gersten *et al*, 2009a; Roberts *et al*, 2004; Roberts *et al*, 2006; Roberts *et al*, 2003; Winkler *et al*, 2012). Over expression of a high number of genes may be due to inflammation and activation of several transcription factors, signaling molecules, and interferon-associated genes, and by presence of virus in the brain. As also shown previously in macaques with acute SIV infection (Roberts *et al*, 2004), increased interferon, innate immunity pathways, and other antiviral responses mediated by macrophages indicated general signs of infection in the brain. For the first time and differently to what previously reported (Roberts *et al*, 2004), however, we found over expression of PARPs, which regulate different aspects of cell metabolism (Bai, 2015) during SIV infection. PARP9 and PARP14 cross-regulate macrophage activation (Iwata *et al*, 2016), while PARP10 and PARP12 are interferon induced genes (Atasheva *et al*, 2014), and have shown antiviral activities such decreasing replication of avian influenza virus (Yu *et al*, 2011) and Zika virus (Li *et al*, 2018), respectively. Overall, PARPs’ activity relationship with host and virus is quite complex, and both pro and antiviral responses have been reported (Kuny and Sullivan, 2016). PARP1-mediated cascade of progression to neurodegeneration and neuroinflammation has been shown in Parkinson’s and Alzheimer’s disease (Martire *et al*, 2015). Yet, PARP1 resulted neither over or under expressed in animals with SIV infection in the frontal cortex, suggesting that its contribution to neuroAIDS may not be significant, despite its known role in HIV suppression by regulating HIV infection and integration (Ha *et al*, 2001; Kameoka *et al*, 2004). On the other hand, four of the 18 PARP genes – PARP9, PARP10, PARP12, PARP14 – were clearly upregulated, suggesting that inflammation may be a byproduct of PARPs activity. Excessive activation of PARPs may cause cell death (Pieper *et al*, 1999), followed by release of cellular components into the CNS, amplification of the immune response, and eventually neurodegeneration.

In summary, we found evidence that PARPs dysregulation could provide new, key indicators of SIV brain infection and neuropathogenesis. Moreover, since PARP inhibitors have shown promising neuroprotective properties (Rumbaugh *et al*, 2008), similar inhibitors may be employed against HIV-related toxicity and inflammation in the brain. Additional statistical studies using a larger number of animals and *in vitro* experiments are needed to determine what is the role of each PARP, and which proteins within PARP-mediated pathways may offer promising candidates as HAND novel therapeutic targets. Nevertheless, our study provides novel insights that may inform drug screening and development efforts aimed at identifying specific antiviral therapies and a new class of potential therapeutic candidates for HAND.

## Conclusions

Our study indicates that PARPs are over-expressed during SIV infection of the brain. PARPs may role in SIV-associated neurodegenerative processes. Inhibition of PARPs may provide an effective novel therapeutic target for HIV-related neuropathology.

## Ethics approval and consent to participate

Procedures on the CD8+ lymphocyte-depleted and naturally progressing cohort were conducted with the approval of New England Regional Primate Center at Harvard (Lamers *et al*, 2015) and University Tulane University’s Institutional Animal Care and Use Committee (Rife *et al*, 2016), respectively.

## Availability of data and materials

The datasets generated during and/or analyzed during the current study are available in the Sequence Read Archive (SRA) with the BioProject PRJNA624871 and will be released by SRA after publication, https://www.ncbi.nlm.nih.gov/sra

## Competing interests

The authors declare that they have no competing interests

## Funding

This work was supported by NIH award R01 NS063897. MS is supported in part by the Stephany W. Holloway University Chair in AIDS Research.

## Authors’ Contributions

CM design of the work, analysis, interpretation of data, wrote the manuscript; ASRM analysis, acquisition of data; JJD analysis, acquisition of data; DJN design of the work, analysis; SNR analysis; KW acquisition of data; BRM revised the manuscript; MC analysis; SM revised the manuscript; MCF revised the manuscript; DMA design of the work; AR creation of pipeline used in the work, interpretation of data, revised the manuscript; KCW interpretation of data, revised the manuscript; MS design of the work, interpretation of data, revised the manuscript. All authors read and approved the final manuscript.

---

**Figure S1.**
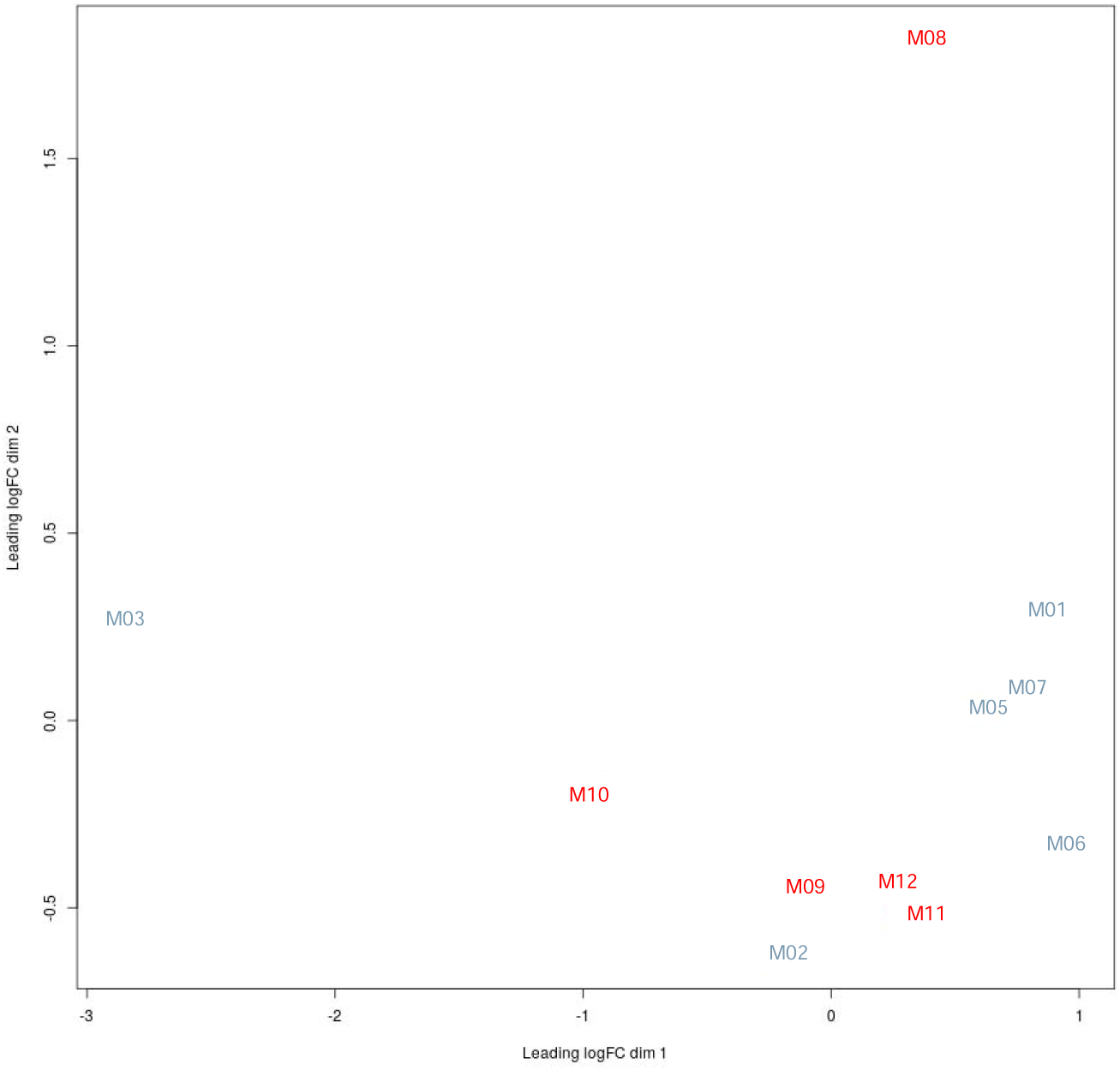
Multi-Dimensional Scaling (MDS) plot for the normalized expression data of depleted versus non depleted SIV-infected animals. Distance based the matrix of FPKM values quantified using RSEM v1.2.31 for all transcripts in all samples of depleted macaques in red, and non-depleted macaques in blue. The plot shows that the groups are overlapping.

## List of abbreviations

cART: combination antiretroviral therapy
HIV: human immunodeficiency virus type 1
HAND: human immunodeficiency virus type 1-associated neurocognitive disorders
SIV: Simian immunodeficiency virus
CNS: central nervous system
neuroAIDS: neurological complications of HIV acquired immunodeficiency syndrome
SIVE: SIV-associated encephalitis
SAIDS: simian AIDS
PARPs: poly(ADP-ribose) polymerases
SGS: single genome sequencing
RNA: ribonucleic acid
mRNA: messenger RNA
qPCR: quantitative polymerase chain reaction
GAPDH: Glyceraldehyde 3-phosphate dehydrogenase
DV: detectable SIV in the brain
NDV: low/undetectable SIV in the brain
DEGs: differentially expressed genes
NPAS: Neuronal PAS Domain Protein
EPSTI: epithelial stromal interaction
SLFN: Schlafen Family Member
MAMU-A: major histocompatibility complex, class I, A (Rhesus monkey)
IFN: interferon
ISGs: type I interferon-stimulated genes
DDX: DExD/H-Box Helicase
RIG1: retinoic acid-inducible gene I
PSMB: Proteasome 20S Subunit Beta
NCF: Neutrophil Cytosolic Factor
STAT: Signal Transducer And Activator Of Transcription
CSF1: Colony Stimulating Factor 1
CSF1R: Colony Stimulating Factor 1 receptor
MNDA: myeloid cell nuclear differentiation antigen
MIF: Macrophage migration inhibitory factor
IFI: Interferon Induced Protein
IRF: Interferon Regulatory Factor
MX1: MX Dynamin Like GTPase 1
OAS1: 2’-5’-Oligoadenylate Synthetase 1
TLR: toll-like receptor
NADPH: Reduced nicotinamide adenine dinucleotide phosphate
C1Q: complement component 1q
C3: complement component 3
TNFSF10: TNF Superfamily Member 10
Th: T helper cell

## Notes

### Competing Interest Statement

The authors have declared no competing interest.

### Summary of Updates

title has changed, added a new figure, moved tables and added more results

