## supplementary material for "Brain tissue transcriptomic analysis of SIV-infected macaques identifies several altered metabolic pathways linked to neuropathogenesis, and Poly (ADP-ribose) polymerases (PARPs) as potential therapeutic targets"

**Supplementary materials**

**Table S1. RNA-seq to *Macaca mulatta* transcriptome.**

| Sample | Total nt | Coverage | Effective bp | Effective % | Effective coverage |
| --- | --- | --- | --- | --- | --- |
| M09 | 33,182,531,269 | 11.73 | 722,689,555 | 25.50% | 45.92 |
| M10 | 27,880,370,047 | 9.85 | 669,288,118 | 23.60% | 41.66 |
| M11 | 29,380,791,916 | 10.39 | 672,744,043 | 23.80% | 43.67 |
| M12 | 31,478,903,107 | 11.13 | 714,766,167 | 25.30% | 44.04 |
| M02 | 32,270,278,512 | 11.41 | 699,306,285 | 24.70% | 46.15 |
| M03 | 4,139,754,570 | 1.46 | 222,551,690 | 7.90% | 18.60 |
| M08 | 4,256,639,530 | 1.51 | 238,088,682 | 8.40% | 17.88 |
| M06 | 40,671,217,226 | 14.38 | 761,736,841 | 26.90% | 53.39 |
| M07 | 24,162,699,505 | 8.54 | 574,793,687 | 20.30% | 42.04 |
| M01 | 33,356,344,328 | 11.79 | 721,917,050 | 25.50% | 46.21 |
| M05 | 30,432,512,038 | 10.76 | 673,555,026 | 23.80% | 45.18 |

nt = nucleotides; bp = base pais; total nt = the total number of nucleotides sequenced, i.e. the number of aligned reads times the length of each read; coverage = the total nucleotides sequenced divided by the size of the transcriptome; effective bp = the number of bases in the transcriptome; effective percent = the percentage of effective bp to the total transcriptome; eff coverage = the average coverage over the effective bp fraction of the transcriptome.

**Table S2. Up- and down-regulated genes and their function related to HIV/SIV.** Differentially expressed genes in frontal cortex of SIV macaques with detectable virus in the brain with log2(FC) >= 1.0 and P-value <= 0.05.

| **Over-expressed Gene** | **Log2(FC)** | **P-value** | **References** | **What it says** |
| --- | --- | --- | --- | --- |
| PATE2 | -2.149 | 0.0132 |  |  |
| GSTT2/GSTT2B | -1.937 | 0.0375 |  |  |
| SPX | -1.548 | 0.00402 |  |  |
| HMCN1 | -1.224 | 0.0331 | (Q. Li et al., 2009) | Found to be downregulated in HIV-1 infection. |
| NPAS4 | -1.187 | 0.00189 |  |  |
| IFI16 | 1.046 | 0.0341 | 1) (Hotter et al., 2019)  2) (Lu et al., 2016) | 1) Has antiretroviral activity against HIV-1 by binding and inhibiting host transcription factor Sp1, which drives viral gene expression 2) SIV infected rhesus macaques in the rectal draining lymph node IFI16 was found to be upregulated |
| FES | 1.097 | 0.0132 |  |  |
| FGD2 | 1.114 | 0.0487 |  |  |
| GPR34 | 1.119 | 0.0188 |  |  |
| HAVCR2 | 1.119 | 0.00646 | (Bosinger et al., 2009) | Genes whose expression correlated strongly with the level of CD8+Ki-67+ T cells in the SIVmac239-infected rhesus macaques may be candidate mediators of the immune activation observed during pathogenic SIV infection, such as HAVCR2 |
| CTSS | 1.119 | 0.0267 | (Borjabad et al., 2011) | Correlated positively with plasma viral load in HIV-1 infected. |
| TNFSF10 | 1.131 | 0.0423 | (Zahoor, Xue, Sato, & Aida, 2015) | HIV-1 Vpr showed to upregulate IFIT1 in monocyte derived macrophages |
| IRF8 | 1.147 | 0.00817 | (D'Antoni et al., 2019) | Intracellular IRF-8 expression was measured in cryopreserved peripheral blood mononuclear cells from chronically HIV-infected individuals on ART and was found to be highest in plasmacytoid dendritic cells |
| CD99 | 1.182 | 0.0375 |  |  |
| CSF1R | 1.183 | 0.0000176 | 1) (Knight, Brill, Queen, Tarwater, & Mankowski, 2018)  2) (Irons, Meinhardt, Allers, Kuroda, & Kim, 2019) | 1) Upregulated in a SIV/macaque model of HIV CNS disease  2) Brain macrophages including perivascular macrophages (PVMs), expressed higher levels of CSF1R compared to microglia. Found significantly increased expression of CSF1R on the infected PVMs and lesional macrophages in the brains of encephalitic macaques with SIV. Moreover, the per cell expression of CSF1R determined by its mean pixel intensity (MPI) correlated positively with the MPI of SIV Gag p28 in SIV-infected PVMs |
| HK2 | 1.192 | 0.0351 |  |  |
| SLC2A5 | 1.205 | 0.0255 | (Winkler, Chaudhuri, & Fox, 2012) | Found to be upregulated in SIV infected monkeys. |
| RPS6KA1 | 1.214 | 0.0179 | (Y. Li, Chan, & Katze, 2007) | Found to be up regulated in HIV-1 infected in pigtail macaque PBMCs. |
| TRIM22 | 1.219 | 0.0153 | (Turrini et al., 2019) | Functions as an antiviral factor by limiting Sp1 transcription in HIV-1 infected CD4+ T cells |
| PML | 1.243 | 0.00000956 | (Lu et al., 2014) | Upregulated in SIV-infected rhesus macaques, infected via rectal mucosa |
| EPHX1 | 1.247 | 0.00187 |  |  |
| TRIM38 | 1.268 | 0.00273 |  |  |
| DOCK8 | 1.28 | 0.00452 |  |  |
| HCLS1 | 1.299 | 0.0267 |  |  |
| HCK | 1.362 | 0.0256 |  |  |
| MYO1F | 1.362 | 0.0324 |  |  |
| PLCG2 | 1.388 | 0.000943 |  |  |
| FAM111A | 1.433 | 0.0269 |  |  |
| VMO1 | 1.445 | 0.00189 |  |  |
| C3AR1 | 1.449 | 0.0171 | 1) (Zahoor et al., 2015)  2) (Nowlin et al., 2018) | 1) Found upregulated by HIV-1 Vpr protein  2) Upregulated in SIV infected intermediate monocytes |
| ALDH3B1 | 1.455 | 0.0124 | (Wu, Sasse, Saksena, & Saksena, 2013) | Found to be upregulated in HIV+ patients on HAART |
| STAT2 | 1.472 | 9.37E-08 | (W. Li, Gorantla, Gendelman, & Poluektova, 2017) | Found to be upregulated in the corpus callosum and hippocampus of HIV-1 infected mice with a humanized brain and immune system |
| TRIM25 | 1.478 | 0.0000509 | (Bosinger et al., 2009) | Viral restriction factor upregulated in SIV-infected sooty mangabays |
| LAPTM5 | 1.493 | 0.000000978 | 1) (Winkler et al., 2012)  2) (Roberts et al., 2003) | 1) Found to be upregulated in SIV encephalitis  2) Upregulated in frontal lobe of SIV encephalitis |
| TLR4 | 1.497 | 0.00404 | 1) (Hernandez, Stevenson, Latz, & Urcuqui-Inchima, 2012)  2) (Mothapo et al., 2017) | 1) An increase of TLR4 expression was observed in monocyte-derived macrophage and PBMCs infected with HIV-1 in vitro and in response to TLR stimulation, compared to the mock. The *ex vivo* analysis indicated increased expression of TLR4 in myeloid dendritic cells  2) Cerebrospinal fluid soluble TLR4 concentrations were higher in SIV-infected macaques with neurological sequelae compared to those without neurological complications. Also, in humans, elevated cerebrospinal fluid soluble TLR4 levels were found in HIV-infected patients with cognitive impairments compared to HIV-infected patients with normal cognition |
| SERPINA1 | 1.5 | 0.0224 | (S. M. Gonzalez et al., 2015) | Upregulated in genital mucosa and gut associated lymphoid tissue in HIV infected individuals |
| PLA2G4C | 1.569 | 0.00641 | (Lu et al., 2014) | Upregulated in SIV-infected rhesus macques, infected via rectal mucosa |
| C3 | 1.57 | 0.00221 | (He et al., 2014) | Previously recognize as a gene with a link to HAND. Upregulated in brain tissues of HIV-infected patients with no ART. |
| S100A6 | 1.574 | 0.0493 |  |  |
| CCR1 | 1.59 | 0.00000435 | (Lu et al., 2014) | Upregulated in SIV-infected rhesus macques, infected via rectal mucosa |
| PTPRC | 1.591 | 0.00744 | (He et al., 2014) | Previously recognize as a gene with a link to HAND. Upregulated in brain tissues of HIV-infected patients with no ART. |
| SP110 | 1.619 | 0.0493 |  |  |
| CD14 | 1.642 | 0.0306 | 1) (Stewart et al., 2020)  2) (R. G. Gonzalez et al., 2018) | 1) In HIV+ patients, greater depressive symptoms were associated with higher soluble CD14. 2) Neuronal injury was found to be correlated with plasma viral levels and infected CD14+CD16+ activated monocytes, known to traffic the virus to the brain, in SIV infected rhesus macaques |
| HMOX1 | 1.649 | 0.00646 | (Gill et al., 2014) | HO-1 protein expression is decreased in the brains of HIV-infected individuals diagnosed with HIV-associated neurocognitive disorders (HAND) and this reduction of HO-1 is associated with CNS viral load and markers of neuroimmune activation, including type I interferon responses |
| ABI3 | 1.659 | 0.0306 |  |  |
| TRIM14 | 1.674 | 0.0000135 |  |  |
| MPEG1 | 1.686 | 0.00061 |  |  |
| HLA-F | 1.704 | 0.00583 | (Tremblay-McLean et al., 2017) | Upregulated in HIV infected T cells |
| UBA7 | 1.711 | 0.000876 |  |  |
| ICAM2 | 1.716 | 0.0239 |  |  |
| BTN3A3 | 1.716 | 0.00000169 | (Gelman et al., 2012) | Found to be upregulated in HIV encephalitis |
| SPI1 | 1.723 | 0.00835 |  |  |
| CTSC | 1.726 | 0.0219 |  |  |
| MNDA | 1.745 | 0.00354 |  |  |
| SP100 | 1.757 | 0.037 | (Lorenz, Misra, & Gabuzda, 2019) | Upregulated expression in HIV+ human monocytes |
| BIN2 | 1.771 | 0.00529 |  |  |
| LCP1 | 1.778 | 0.0078 |  |  |
| OASL | 1.786 | 0.0471 | (Sanfilippo et al., 2018) | Upregulated in macaques with SIV encephalitis and patients with HIV associated neurocognitive disorders |
| UBE2L6 | 1.796 | 0.000628 | (Siangphoe & Archer, 2015) | Upregulated in brain from HIV-associated neurocognitive disorders, HIV encephalitis and HIV-infected patients versus controls. |
| RNF213 | 1.807 | 0.0000344 |  |  |
| S100A11 | 1.809 | 0.0211 |  |  |
| TM4SF1 | 1.825 | 0.0226 |  |  |
| **PARP9** | 1.837 | 0.0049 |  |  |
| TMIGD3 | 1.866 | 0.026 |  |  |
| IRF9 | 1.871 | 0.00000169 | (W. Li et al., 2017) | Upregulated in brain tissues from HIV-infected mice |
| APOBEC3G | 1.875 | 0.0486 | (De Scheerder et al., 2020) | Upregulated in HIV-1 seroconverters, ART-naïve acutely infected HIV-1 patients |
| **PARP12** | 1.89 | 0.00067 |  |  |
| **PARP10** | 1.907 | 0.00000519 |  |  |
| BIRC5 | 1.932 | 0.0149 |  |  |
| CD37 | 1.943 | 0.0000246 | 1) (Matheson et al., 2015)  2) (Haller et al., 2014)  3) (Mohan et al., 2013) | 1) Downregulated during HIV infection  2) Downregulated by HIV-1 Nef  3) Upregulated in intestinal epithelium of Indian-origin rhesus macaques infected with SIV |
| TLR3 | 1.946 | 0.0249 | (Bhargavan & Kanmogne, 2018) | Upregulated in HIV+ patients in brain tissues and blood vessels, even higher levels of expression in patients with HAND |
| STAT1 | 1.95 | 0.000000746 | (Gelman et al., 2012) | Its mRNA showed significant correlation with brain viral load from HIV-1 infected subjects with no substantial neurocognitive impairment (NCI), Infected with substantial NCI without HIV encephalitis (HIVE), and Infected with substantial NCI and HIVE |
| GFAP | 1.951 | 0.0105 | 1) (Guha et al., 2019)  2)(Wachter, Eiden, Naumann, Depboylu, & Weihe, 2016) | 1) Increased expression in cerebrospinal fluid from subjects with HAND versus without. 2) SIV-infected AIDS-diseased rhesus macaques showed increased expression of GFAP in astrocytes of subcortical white matter |
| CYBB | 1.958 | 0.0219 |  |  |
| CD48 | 1.972 | 0.0179 |  |  |
| OAS3 | 2.003 | 0.000000131 | (Lu et al., 2014) | Upregulated in SIV-infected rhesus macques, infected via rectal mucosa |
| NMI | 2.017 | 0.0127 |  |  |
| C1QB | 2.036 | 0.00109 | (Vlasova-St Louis, Chang, Shahid, French, & Bohjanen, 2018) | Upregulated in HIV-infected subjects with cryptococcal meningitis with ART and identified as an early and late development of cryptococcosis-associated immune reconstitution inflammatory syndrome |
| APOL2 | 2.05 | 0.00744 |  |  |
| HLA-A | 2.054 | 0.0238 |  |  |
| HLA-E | 2.065 | 0.0000137 |  |  |
| PARVG | 2.076 | 0.0269 |  |  |
| IFI6 | 2.08 | 0.0392 | (W. Li et al., 2017) | Found to be upregulated in mice infected with HIV-1 |
| TYMS | 2.097 | 0.0331 |  |  |
| IFI35 | 2.097 | 0.0000199 | 1) (Lu et al., 2014)  2) (Lorenz et al., 2019) | 1) Upregulated in SIV-infected rhesus macques, infected via rectal mucosa  2) Upregulated in monocytes from HIV+ smokers compared with HIV− non-smokers, also associated with depressive symptoms in logistic regression models adjusted for HIV status and smoking |
| BTN3A1 | 2.122 | 0.000000214 | (Gelman et al., 2012) | Its mRNA showed significant correlation with brain viral load from HIV-1 infected subjects with no substantial neurocognitive impairment (NCI), Infected with substantial NCI without HIV encephalitis (HIVE), and Infected with substantial NCI and HIVE |
| BTN2A1 | 2.123 | 0.0227 |  |  |
| ARHGAP15 | 2.169 | 0.0129 |  |  |
| DDX60L | 2.17 | 0.0000344 |  |  |
| ALDH3A1 | 2.18 | 0.0181 |  |  |
| MICB | 2.194 | 0.00109 | 1) (Omi et al., 2014)  2) (Averdam et al., 2007) | Upregulated in CCR5‐expressing HIV‐1 infected CD4+ NKT cells  2) Mamu-MIC genotyping of DNA of a cohort of 68 experimentally SIV-infected rhesus macaques revealed no significant association of either of the two Mamu-MICA or MICB allelic lineages with differences in progression to AIDS-like symptoms |
| TRIM21 | 2.194 | 0.000449 | (Ross, Roodgar, & Smith, 2015) | Showed to be a strong candidate gene for SIVmac resistance in Chinese rhesus macaques |
| TFEC | 2.21 | 0.0325 |  |  |
| CCR5 | 2.215 | 0.00314 | 1) (Kim et al., 2018)  2) (M. Zhou et al., 2016)  3) (Delery et al., 2019) | 1) It has been suggested a critical role for glial CCR5 in mediating neurotoxic effects of HIV-1 Tat and morphine interactions on neurons  2) CCR5 plays an important role in neuroplasticity, learning and memory, and it has a role in the cognitive deficits caused by HIV  3) Lower numbers of CCR5-positive cells in the brain, combined with a less leaky blood-brain barrier, may be responsible for the decreased virus infection in the brain and consequently the absence of encephalitis in newborn macaques infected with SIV |
| UNC93B1 | 2.249 | 0.0000276 |  |  |
| HRASLS2 | 2.272 | 0.00927 | (Fisher et al., 2018) | Up regulated in SIV-infected infant macaques. |
| ADA2 | 2.276 | 0.0362 |  |  |
| DDX58 | 2.299 | 0.00000169 | (Zahoor et al., 2014) | HIV-1 Vpr showed to upregulate DDX58 in monocyte derived macrophages |
| SAMD9L | 2.304 | 0.0000486 | (Hogerkorp, Nishimura, Song, Martin, & Roederer, 2011) | Up regulated in T cells from rhesus macaques with acute SIVmac239 infection |
| DHX58 | 2.336 | 0.000000126 | (Lenz et al., 2015) | The expression level of DHX58 has showed an association with HIV-1 viral load. |
| SLFN13 | 2.35 | 0.0037 | (Lu et al., 2014) | Upregulated in SIV-infected rhesus macques, infected via rectal mucosa |
| B2M | 2.363 | 0.00000104 | (Lu et al., 2014) | Upregulated in SIV-infected rhesus macques, infected via rectal mucosa |
| C1QC | 2.368 | 0.00332 | (Rempel et al., 2013) | Upregulated in HIV-suppressed coinfected subjects (HIV – HCV) which also correlated with global deficit score thereby linking an expression profile with poorer cognition |
| FCGR3A/FCGR3B | 2.382 | 0.00564 | (Lu et al., 2014) | Upregulated in SIV-infected rhesus macques, infected via rectal mucosa |
| HERC6 | 2.43 | 1.83E-08 | 1) (Lu et al., 2014)  2) (W. Li et al., 2017) | 1) Upregulated in SIV-infected rhesus macques, infected via rectal mucosa  2) Upregulated in the hippocampus and corpus callosum in HIV-infected humanized mice |
| HELZ2 | 2.461 | 0.000000149 |  |  |
| OAS1 | 2.509 | 0.0000509 | 1) (Lu et al., 2014)  2) (Wie et al., 2013) | 1) Upregulated in SIV-infected rhesus macques, infected via rectal mucosa  2) Downregulated in macrophages treated with IFNs and then infected with HIV |
| LGALS3BP | 2.521 | 0.00000698 | 1) (Lodermeyer et al., 2018)  2) (Bachtel et al., 2019) | 1) Upregulated in HIV infection  2)Upregulated in ART-suppressed individuals in memory CD4+ T cells |
| IFIH1 | 2.559 | 0.00000116 | 1) (Ellegard et al., 2014)  2) (Sandler et al., 2014)  3) (Lederer et al., 2009) | 1) Upregulated in immature dendritic cells exposed to free HIV-1 for 24 h  2) Downregulated in rhesus macaques with SIV infection  3) Upregulated in SIV-infected African green monkeys |
| HLA-B | 2.568 | 0.000943 |  |  |
| ZC3HAV1 | 2.573 | 0.0000354 | 1) (Wang, Huang, Jiang, & Sun, 2011)  2) (Bosinger et al., 2009) | 1) Upregulated in HIV encephalitis  2) SIV-infected sooty mangabays exhibit increased expression of the host-restriction genes TRIM22 and ZC3HAV1/ZAP during the acute phase |
| OAS2 | 2.618 | 0.00000169 | 1) (Lu et al., 2014)  2) (Chang et al., 2013) | 1) Upregulated in SIV-infected rhesus macques, infected via rectal mucosa  2) Its expression levels were significantly associated with the level of viral replication or CD4+ T-cell counts in treatment-naive HIV-1–infected individuals |
| IFI44 | 2.62 | 0.00000169 | 1) (Rotger et al., 2011)  2) (Hubbard et al., 2012)  3) (George et al., 2014) | 1) Upregulated in rapid HIV-1 progressors versus viremic nonprogressors (criteria in methods)  2) Showed increased expression that was inversely correlated with HIV load decline in HIV-1 monoinfected patients with peginterferon alfa-2a treatment  3) Upregulated in the circulating CD8+ T cells of rhesus macaque at 3 weeks post infection with SIVmac251 |
| CD74 | 2.628 | 0.00281 | (Trifone et al., 2018) | Upregulated by HIV Nef and Vpu and the magnitude of this upregulation correlates with the level of immune activation in the HIV+ subjects |
| CYTIP | 2.64 | 0.00301 |  |  |
| XAF1 | 2.641 | 0.00000196 | 1) (Wie et al., 2013)  2) (Bosinger et al., 2009) | 1) Downregulated in macrophages treated with IFNs and then infected with HIV  2) SIV-infected sooty mangabays exhibited lower expression levels of XAF1 during the acute phase of SIV infection |
| IRF7 | 2.656 | 0.000759 | 1) (Zahoor et al., 2014)  2) (Chang et al., 2013)  3) (Alammar, Gama, & Clements, 2011) | 1) HIV-1 Vpr showed to upregulate IRF7 in monocyte derived macrophages  2) Its expression levels were significantly associated with the level of viral replication or CD4+ T-cell counts in treatment-naive HIV-1–infected individuals  3) Expression of IRF7 mRNA in the brain during acute infection was upregulated in pigtailed macaques e dual inoculated with an immunosuppressive swarm of SIV (SIV/DeltaB670) and a neurovirulent clone (SIV/17EFr) |
| PLAC8 | 2.723 | 0.0000943 | (Sabado et al., 2010) | Upregulated in myeloid dendritic cells from HIV infected subjects |
| **PARP14** | 2.741 | 0.0000036 |  |  |
| HLA-DRB5 | 2.775 | 0.0315 |  |  |
| LGALS9B | 2.821 | 0.00109 |  |  |
| TLR8 | 2.866 | 0.0209 | (Cervantes et al., 2016) | Found to be upregulated in oral epithelial cells in HIV-1 infeed subjects |
| IFIT1 | 2.885 | 2.46E-10 | 1) (Zahoor et al., 2014)  2) (Lu et al., 2014) | 1) HIV-1 Vpr showed to upregulate IFIT1 in monocyte derived macrophages  2) Upregulated in SIV-infected rhesus macques, infected via rectal mucosa |
| IFIT2 | 2.886 | 1.83E-08 | 1) (Zahoor et al., 2014)  2) (Lu et al., 2014) | 1) HIV-1 Vpr showed to upregulate IFIT2 in monocyte derived macrophages  2) Upregulated in SIV-infected rhesus macques, infected via rectal mucosa |
| MX1 | 2.9 | 0.00000394 | 1) (Rotger et al., 2011)  2) (Zahoor et al., 2014)  3) (Lu et al., 2014) | 1) Upregulated in rapid HIV-1 progressors versus viremic nonprogressors (criteria in methods)  2) HIV-1 Vpr showed to upregulate MX1 in monocyte derived macrophages  3) Upregulated in SIV-infected rhesus macques, infected via rectal mucosa |
| IFI44L | 2.96 | 6.85E-08 | 1) (Rotger et al., 2011)  2) (Zahoor et al., 2014)  3) (Lu et al., 2014) | 1) Upregulated in rapid HIV-1 progressors versus viremic nonprogressors (criteria in methods)  2) HIV-1 Vpr showed to upregulate IFI44L in monocyte derived macrophages  3) Upregulated in SIV-infected rhesus macques, infected via rectal mucosa |
| HLA-DRA | 2.963 | 0.0186 | (Maingat et al., 2011) | Found to be upregulated in HIV+ infected people from duodenal samples |
| EPSTI1 | 2.966 | 0.00452 | (Rotger et al., 2011) | 1) Upregulated in rapid HIV-1 progressors versus viremic nonprogressors (criteria in methods) |
| EFCAB11 | 2.986 | 0.00584 |  |  |
| TLR2 | 2.988 | 0.00183 | 1) (Hernandez et al., 2012)  2) (Heggelund et al., 2004)  3) (Mothapo et al., 2017) | 1) An increase of TLR2 expression was observed in monocyte-derived macrophage and PBMCs infected with HIV-1 in vitro and in response to TLR stimulation, compared to the mock. The ex vivo analysis indicated increased expression of TLR2 in myeloid dendritic cells, but only of TLR2 in monocytes obtained from HIV-1-infected patients, compared to healthy subjects. Remarkably, the expression was higher in cells from patients who do not use HAART  2) Isolated monocytes from HIV-infected patients displayed enhanced expression of TLR2 but not TLR4, that TLR2 expression on the surface of monocytes was significantly increased upon stimulation of HIV type 1 envelope protein gp120, and that TLR2 stimulation in HIV-infected patients induced increased viral replication and TNF-α response  3) Cerebrospinal fluid soluble TLR2 concentrations were higher in SIV-infected macaques with neurological sequelae compared to those without neurological complications. |
| RUNX1 | 2.993 | 0.0222 |  |  |
| SIGLEC1 | 3.011 | 0.00817 | 1) (Zhang et al., 2018)  2) (Jaroenpool et al., 2007) | 1) Was found to be downregulated in untreated HIV-1 infected individuals when compared to treated naïve patients with viremia  2) Monocytes from SIV infected disease resistant sooty mangabeys showed a decrease in SIGLEC1 expression |
| PSMB9 | 3.038 | 0.00000462 | 1) (Lu et al., 2014)  2) (Winkler et al., 2012) | 1) Upregulated in SIV-infected rhesus macaques, intra-rectally inoculated.  2) Upregulated in SIV encephalitis |
| BST2 | 3.093 | 0.000943 | 1) (Homann, Smith, Little, Richman, & Guatelli, 2011)  2) (Mussil, Javed, Topfer, Sauermann, & Sopper, 2015) | 1) The expression of BST-2 was increased on mononuclear leukocytes, including CD4-positive T lymphocytes from HIV-positive patients, compared to that on cells of uninfected controls. The expression of BST-2 was highest during acute infection and decreased to levels similar to those of uninfected individuals after ART  2) When compared to pre-infection levels, BST2 expression was increased in PBMC, purified CD4(+) lymphocytes and CD14(+) monocytes of SIV-infected animals. |
| C1orf162 | 3.12 | 0.000316 |  |  |
| TMPRSS2 | 3.121 | 0.00000169 |  |  |
| CD300A | 3.129 | 0.0362 | (Vitalle et al., 2020) | Overexpressed in CD4+ T cells of HIV+ patients |
| RARRES3 | 3.194 | 0.0165 |  |  |
| ODF3B | 3.207 | 0.000404 |  |  |
| ZBP1 | 3.269 | 0.00881 | (Y. Zhou, Rong, Lu, Pan, & Liang, 2008) | IMP1 interacts with HIV-1 Gag protein and is able to block the formation of infectious HIV-1 particles |
| NLRC5 | 3.303 | 0.0000462 | 1) (Shiau et al., 2019)  2) (Periyasamy, Thangaraj, Bendi, & Buch, 2019) | 1) The top hypomethylated site found in a promoter region was in NLRC5, in HIV-infected subjects, which encodes a transcription factor that regulates major histocompatibility complex (MHC) class I molecule expression  2) Exposure of mouse microglia to HIV-1 Tat resulted both in a dose- and time-dependent upregulation of miRNA-34a, with concomitant downregulation of NLRC5 |
| PSMB8 | 3.303 | 0.00258 | 1) (Gelman et al., 2012)  2) (Yang et al., 2015) | 1) Patients infected with HIV-1 with substantial neurocognitive impairment without HIV encephalitis showed upregulated levels of PSMB8. 2) PSMB8 was upregulated in HIV infection   PSMB8 was upregulated in SIV infected rhesus macaques |
| IFIT3 | 3.312 | 7.15E-08 | 1) (Zahoor et al., 2014)  2) (Nasr et al., 2017)  3) (Ellegard et al., 2014) | 1) HIV-1 Vpr protein upregulates genes involved in the innate response, including IFIT3  2) IFIT3 inhibited HIV production, measured as extracellular infectious virus  3) Immature dendritic cells exposed to free HIV-1 exhibited high mRNA levels of IFIT3. Quantitative proteomics showed upregulation of IFIT3 in immature dendritic cells after a 24 h exposure to free HIV. |
| NCF1 | 3.365 | 0.00528 | (Cobos Jimenez et al., 2015) | NCF1 was found to be one of the genes with antiviral characteristics that was upregulated in macrophages stimulated with IFNγ and TNFα (where [HIV-1 infection](https://www.sciencedirect.com/topics/medicine-and-dentistry/human-immunodeficiency-virus-1-infection) is restricted at a post-integration level) when compared to non-stimulated macrophages |
| C4A/C4B | 3.366 | 0.00666 |  |  |
| MX2 | 3.387 | 7.08E-11 | (Kane et al., 2013) | MX2 is an effector of the anti-HIV-1 activity of type-I IFN. It is suggested that MX2 inhibits HIV-1 infection by inhibiting capsid-dependent nuclear import of subviral complexes |
| IFI27 | 3.553 | 0.00332 | (Zahoor et al., 2015) | Found to be upregulated by HIV-1 Vpr in human dendritic cells |
| HERC5 | 3.596 | 8.88E-09 | 1) (Woods et al., 2011)  2) (Paparisto et al., 2018) | 1) Has found to block an early stage of retroviral Gag particle assembly in HIV-1.  2) Coelacanth HERC5 inhibited SIV but not HIV-1 particle production |
| USP41 | 3.606 | 0.0000509 |  |  |
| SECTM1 | 3.678 | 0.0179 |  |  |
| TYMP | 3.917 | 1.92E-10 | (Williams, Ipser, Stein, Joska, & Naude, 2019) | TYMP is one of the dysregulated neuro-immune markers that has previously been linked to HIV-associated neurocognitive impairment. This marker was found to be significantly higher in HIV+ when compared to HIV- participants. |
| DDX60 | 3.958 | 1.28E-11 | 1) (Lu et al., 2014)  2) (Lu et al., 2016) | 1) It is a viral RNA pattern recognition gene that was found to be upregulated at 6 and 10 dpi after recta SIV exposure in rhesus macaques. DDX60 has also been reported to be an antiviral factor that interacts with RIG-I, MDA5 and LGP2 and regulates RIG-I- and MDA5-mediated type I IFN activation after viral infection  2)  At 6 dpi, RIG-I, melanoma differentiation-associated protein 5 (MDA5), LGP2 and DDX60, all involved in double-stranded RNA (dsRNA) recognition, were upregulated, and genes encoding these proteins were further upregulated at 10 dpi |
| ISG15 | 4.125 | 0.0000344 | 1) (Zahoor et al., 2014)  2) (Kunzi & Pitha, 1996) | 1) HIV-1 Vpr protein (involved in HIV-1 replication and pathogenesis) causes ISG15 (ISGs that can inhibit virus replication) upregulation in monocyte-derived macrophages. Because of this ISG induction by Vpr, macrophages are relatively resistant to Vpr-induced cell death. This suggests a potential role for these genes in host defense against HIV-1 replication and infection  2) A sustained induction of IGS15 may confer a higher therapeutic index to IFN-ω in controlling HIV infection |
| OR1L6 | 4.695 | 0.0404 |  |  |
| ISG20 | 4.701 | 0.00363 | 1) (Zahoor et al., 2014) | 1) HIV-1 Vpr protein (involved in HIV-1 replication and pathogenesis) causes ISG20 (ISGs that can inhibit virus replication) upregulation in monocyte-derived macrophages. Because of this ISG induction by Vpr, macrophages are relatively resistant to Vpr-induced cell death. This suggests a potential role for these genes in host defense against HIV-1 replication and infection |
| IL21R | 4.795 | 0.00000914 | 1)(Pallikkuth, Parmigiani, & Pahwa, 2012)  2) (Bortell, Morsey, Basova, Fox, & Marcondes, 2015) | 1) Up-regulated IL-21R has been described in B cells from HIV-1 infected patients compared to control patients, with the highest levels in non-treated patients.  2) Expression of IL21 was significantly decreased by SIV infection while IL21R levels were increased in macaques. |
| CXCL2 | 5.767 | 0.00409 | (Chen, Yi, Zhang, Klotman, & Chen, 2016) | Found to be upregulated in renal tube epithelial cells infected with HIV-1. |

**Table S3. Predicted enrichment in pathways and genes.** -log(p-value)= significance of overlap of the observed genes and the genes in the pathway; z-score = z-score of the pathway is based on comparison between the direction of the observed genes compared to direction of those same genes in the active state of the pathway.

| **Pathways** | **-log(p-value)** | **z-score** | **Observed gene in pathway** |
| --- | --- | --- | --- |
| Activation of IRF by Cytosolic Pattern Recognition Receptors | 1.06E+01 | 1.9 | DDX58, DHX58, IFIH1, IFIT2, IRF7, IRF9, ISG15, STAT1, STAT2, ZBP1 |
| Calcium-induced T Lymphocyte Apoptosis | 2.90E+00 | 2 | HLA-A, HLA-B, HLA-DRA, HLA-DRB5 |
| Complement System | 5.21E+00 | 2 | C1QB, C1QC, C3, C3AR1, C4A/C4B |
| Death Receptor Signaling | 3.32E+00 | 2.2 | PARP10, PARP12, PARP14, PARP9, TNFSF10 |
| Dendritic Cell Maturation | 8.80E+00 | 3.4 | B2M, FCGR3A/FCGR3B, HLA-A, HLA-B, HLA-DRA, HLA-DRB5, IRF8, PLCG2, STAT1, STAT2, TLR2, TLR3, TLR4 |
| Fcy Receptor-mediated Phagocytosis in Macrophages and Monocytes | 2.34E+00 | 2 | FCGR3A/FCGR3B, HCK, HMOX1, NCF1 |
| iCOS-iCOSL Signaling in T Helper Cells | 2.70E+00 | 2 | HLA-A, HLA-B, HLA-DRA, HLA-DRB5, PTPRC |
| Interferon Signaling | 1.51E+01 | 3.2 | IFI35, IFI6, IFIT1, IFIT3, IRF9, ISG15, MX1, OAS1, PSMB8, STAT1, STAT2 |
| MIF Regulation of Innate Immunity | 3.65E+00 | 2 | CD14, CD74, PLA2G4C, TLR4 |
| MIF-mediated Glucocorticoid Regulation | 4.02E+00 | 2 | CD14, CD74, PLA2G4C, TLR4 |
| Neuroinflammation Signaling Pathway | 1.00E+01 | 3.6 | B2M, BIRC5, CSF1R, CYBB, HLA-A, HLA-B, HLA-DRA, HLA-DRB5, HMOX1, IRF7, PLA2G4C, PLCG2, STAT1, TLR2, TLR3, TLR4, TLR8 |
| NF-β Signaling | 1.94E+00 | 2.2 | PLCG2, TLR2, TLR3, TLR4, TLR8 |
| Phospholipases | 2.92E+00 | 2 | HMOX1, PLA2G4C, PLCG2, RARRES3 |
| PI3K Signaling in B Lymphocytes | 1.77E+00 | 2 | C3, PLCG2, PTPRC, TLR4 |
| PKC-θ Signaling in T Lymphocytes | 2.13E+00 | 2.2 | HLA-A, HLA-B, HLA-DRA, HLA-DRB5, PLCG2 |
| Production of Nitric Oxide and Reactive Oxygen Species in Macrophages | 4.80E+00 | 2.8 | CYBB, IRF8, NCF1, PLCG2, SERPINA1, SPI1, STAT1, TLR2, TLR4 |
| Role of NFAT in Regulation of the Immune Response | 2.55E+00 | 2.4 | FCGR3A/FCGR3B, HLA-A, HLA-B, HLA-DRA, HLA-DRB5, PLCG2 |
| Role of RIG1-like Receptors in Antiviral Innate Immunity | 4.83E+00 | 1.3 | DDX58, DHX58, IFIH1, IRF7, TRIM25 |
| T Cell Exhaustion Signaling Pathway | 6.92E+00 | 1.3 | HAVCR2, HLA-A, HLA-B, HLA-DRA, HLA-DRB5, HLA-E, HLA-F, IRF9, PLCG2, STAT1, STAT2 |
| Tec Kinase Signaling | 2.74E+00 | 2.2 | HCK, PLCG2, STAT1, STAT2, TLR4, TNFSF10 |
| Th1 Pathway | 4.25E+00 | 2.2 | CCR5, HAVCR2, HLA-A, HLA-B, HLA-DRA, HLA-DRB5, STAT1 |
| Toll-like Receptor Signaling | 3.68E+00 | 2 | CD14, TLR3, TLR4 |
| TREM1 Signaling | 4.80E+00 | 2.4 | NLRC5, PLCG2, TLR3, TLR4 |

**Table S4. Extended table of enriched pathways and genes. (excel file)**

**Table S5. Expression of PARPs genes.** Differentially expressed genes in frontal cortex of SIV macaques with detectable virus in the brain with log2(FC) >= 1.0 and P-value <= 0.05.

| **PARPs** | **Log2(FC)** | **P-value** | **SE** | **BaseAvg** |
| --- | --- | --- | --- | --- |
| PARP1 | 0.046819 | 0.940203 | 0.099247 | 882.7174 |
| PARP2 | 0.158777 | 0.796313 | 0.145152 | 221.6855 |
| PARP3 | 1.045341 | 0.083354 | 0.326347 | 74.82065 |
| PARP4 | 0.280497 | 0.74453 | 0.219763 | 408.1099 |
| PARP6 | -0.10317 | 0.822764 | 0.103833 | 613.2247 |
| PARP8 | -0.04821 | 0.973672 | 0.229319 | 75.13621 |
| PARP9 | 1.836935 | 0.004898 | 0.438954 | 113.5548 |
| PARP10 | 1.907221 | 5.19E-06 | 0.333092 | 326.299 |
| PARP11 | -0.43835 | 0.775054 | 0.373313 | 17.8861 |
| PARP12 | 1.890447 | 0.00067 | 0.402328 | 23.23585 |
| PARP14 | 2.74146 | 3.6E-06 | 0.471785 | 93.30319 |
| PARP15 | 1.992009 | 0.055911 | 0.59088 | 14.05644 |
| PARP16 | 0.506024 | 0.428362 | 0.235798 | 66.26066 |

**Table S2 References**

Alammar, L., Gama, L., & Clements, J. E. (2011). Simian immunodeficiency virus infection in the brain and lung leads to differential type I IFN signaling during acute infection. *J Immunol, 186*(7), 4008-4018. doi:10.4049/jimmunol.1003757

Averdam, A., Seelke, S., Grutzner, I., Rosner, C., Roos, C., Westphal, N., . . . Walter, L. (2007). Genotyping and segregation analyses indicate the presence of only two functional MIC genes in rhesus macaques. *Immunogenetics, 59*(3), 247-251. doi:10.1007/s00251-006-0187-1

Bachtel, N. D., Beckerle, G. A., Mota, T. M., Rougvie, M. M., Raposo, R. A. S., Jones, R. B., . . . Apps, R. (2019). Short Communication: Expression of Host Restriction Factors by Memory CD4+ T Cells Differs Between Healthy Donors and HIV-1-Infected Individuals with Effective Antiretroviral Therapy. *AIDS Res Hum Retroviruses, 35*(1), 108-111. doi:10.1089/aid.2018.0162

Bhargavan, B., & Kanmogne, G. D. (2018). Toll-Like Receptor-3 Mediates HIV-1-Induced Interleukin-6 Expression in the Human Brain Endothelium via TAK1 and JNK Pathways: Implications for Viral Neuropathogenesis. *Mol Neurobiol, 55*(7), 5976-5992. doi:10.1007/s12035-017-0816-8

Borjabad, A., Morgello, S., Chao, W., Kim, S. Y., Brooks, A. I., Murray, J., . . . Volsky, D. J. (2011). Significant effects of antiretroviral therapy on global gene expression in brain tissues of patients with HIV-1-associated neurocognitive disorders. *PLoS Pathog, 7*(9), e1002213. doi:10.1371/journal.ppat.1002213

Bortell, N., Morsey, B., Basova, L., Fox, H. S., & Marcondes, M. C. (2015). Phenotypic changes in the brain of SIV-infected macaques exposed to methamphetamine parallel macrophage activation patterns induced by the common gamma-chain cytokine system. *Front Microbiol, 6*, 900. doi:10.3389/fmicb.2015.00900

Bosinger, S. E., Li, Q., Gordon, S. N., Klatt, N. R., Duan, L., Xu, L., . . . Kelvin, D. J. (2009). Global genomic analysis reveals rapid control of a robust innate response in SIV-infected sooty mangabeys. *J Clin Invest, 119*(12), 3556-3572. doi:10.1172/jci40115

Cervantes, C. A., Oliveira, L. M., Manfrere, K. C., Lima, J. F., Pereira, N. Z., Duarte, A. J., & Sato, M. N. (2016). Antiviral factors and type I/III interferon expression associated with regulatory factors in the oral epithelial cells from HIV-1-serodiscordant couples. *Sci Rep, 6*, 25875. doi:10.1038/srep25875

Chang, J. J., Woods, M., Lindsay, R. J., Doyle, E. H., Griesbeck, M., Chan, E. S., . . . Altfeld, M. (2013). Higher expression of several interferon-stimulated genes in HIV-1-infected females after adjusting for the level of viral replication. *J Infect Dis, 208*(5), 830-838. doi:10.1093/infdis/jit262

Chen, P., Yi, Z., Zhang, W., Klotman, M. E., & Chen, B. K. (2016). HIV infection-induced transcriptional program in renal tubular epithelial cells activates a CXCR2-driven CD4+ T-cell chemotactic response. *Aids, 30*(12), 1877-1888. doi:10.1097/qad.0000000000001153

Cobos Jimenez, V., Martinez, F. O., Booiman, T., van Dort, K. A., van de Klundert, M. A., Gordon, S., . . . Kootstra, N. A. (2015). G3BP1 restricts HIV-1 replication in macrophages and T-cells by sequestering viral RNA. *Virology, 486*, 94-104. doi:10.1016/j.virol.2015.09.007

D'Antoni, M. L., Kallianpur, K. J., Premeaux, T. A., Corley, M. J., Fujita, T., Laws, E. I., . . . Ndhlovu, L. C. (2019). Lower Interferon Regulatory Factor-8 Expression in Peripheral Myeloid Cells Tracks With Adverse Central Nervous System Outcomes in Treated HIV Infection. *Front Immunol, 10*, 2789. doi:10.3389/fimmu.2019.02789

De Scheerder, M. A., Van Hecke, C., Zetterberg, H., Fuchs, D., De Langhe, N., Rutsaert, S., . . . Vandekerckhove, L. (2020). Evaluating predictive markers for viral rebound and safety assessment in blood and lumbar fluid during HIV-1 treatment interruption. *J Antimicrob Chemother, 75*(5), 1311-1320. doi:10.1093/jac/dkaa003

Delery, E., Bohannon, D. G., Irons, D. L., Allers, C., Sugimoto, C., Cai, Y., . . . Kim, W. K. (2019). Lack of susceptibility in neonatally infected rhesus macaques to simian immunodeficiency virus-induced encephalitis. *J Neurovirol, 25*(4), 578-588. doi:10.1007/s13365-019-00755-w

Ellegard, R., Crisci, E., Burgener, A., Sjowall, C., Birse, K., Westmacott, G., . . . Larsson, M. (2014). Complement opsonization of HIV-1 results in decreased antiviral and inflammatory responses in immature dendritic cells via CR3. *J Immunol, 193*(9), 4590-4601. doi:10.4049/jimmunol.1401781

Fisher, B. S., Green, R. R., Brown, R. R., Wood, M. P., Hensley-McBain, T., Fisher, C., . . . Sodora, D. L. (2018). Liver macrophage-associated inflammation correlates with SIV burden and is substantially reduced following cART. *PLoS Pathog, 14*(2), e1006871. doi:10.1371/journal.ppat.1006871

Gelman, B. B., Chen, T., Lisinicchia, J. G., Soukup, V. M., Carmical, J. R., Starkey, J. M., . . . Morgello, S. (2012). The National NeuroAIDS Tissue Consortium brain gene array: two types of HIV-associated neurocognitive impairment. *PLoS One, 7*(9), e46178. doi:10.1371/journal.pone.0046178

George, M. D., Hu, W., Billingsley, J. M., Reeves, R. K., Sankaran-Walters, S., Johnson, R. P., & Dandekar, S. (2014). Transcriptional profiling of peripheral CD8+T cell responses to SIVDeltanef and SIVmac251 challenge reveals a link between protective immunity and induction of systemic immunoregulatory mechanisms. *Virology, 468-470*, 581-591. doi:10.1016/j.virol.2014.09.013

Gill, A. J., Kovacsics, C. E., Cross, S. A., Vance, P. J., Kolson, L. L., Jordan-Sciutto, K. L., . . . Kolson, D. L. (2014). Heme oxygenase-1 deficiency accompanies neuropathogenesis of HIV-associated neurocognitive disorders. *J Clin Invest, 124*(10), 4459-4472. doi:10.1172/jci72279

Gonzalez, R. G., Fell, R., He, J., Campbell, J., Burdo, T. H., Autissier, P., . . . Ratai, E. M. (2018). Temporal/compartmental changes in viral RNA and neuronal injury in a primate model of NeuroAIDS. *PLoS One, 13*(5), e0196949. doi:10.1371/journal.pone.0196949

Gonzalez, S. M., Taborda, N. A., Feria, M. G., Arcia, D., Aguilar-Jimenez, W., Zapata, W., & Rugeles, M. T. (2015). High Expression of Antiviral Proteins in Mucosa from Individuals Exhibiting Resistance to Human Immunodeficiency Virus. *PLoS One, 10*(6), e0131139. doi:10.1371/journal.pone.0131139

Guha, D., Lorenz, D. R., Misra, V., Chettimada, S., Morgello, S., & Gabuzda, D. (2019). Proteomic analysis of cerebrospinal fluid extracellular vesicles reveals synaptic injury, inflammation, and stress response markers in HIV patients with cognitive impairment. *J Neuroinflammation, 16*(1), 254. doi:10.1186/s12974-019-1617-y

Haller, C., Muller, B., Fritz, J. V., Lamas-Murua, M., Stolp, B., Pujol, F. M., . . . Fackler, O. T. (2014). HIV-1 Nef and Vpu are functionally redundant broad-spectrum modulators of cell surface receptors, including tetraspanins. *J Virol, 88*(24), 14241-14257. doi:10.1128/jvi.02333-14

He, H., Sharer, L. R., Chao, W., Gu, C. J., Borjabad, A., Hadas, E., . . . Volsky, D. J. (2014). Enhanced human immunodeficiency virus Type 1 expression and neuropathogenesis in knockout mice lacking Type I interferon responses. *J Neuropathol Exp Neurol, 73*(1), 59-71. doi:10.1097/nen.0000000000000026

Heggelund, L., Muller, F., Lien, E., Yndestad, A., Ueland, T., Kristiansen, K. I., . . . Froland, S. S. (2004). Increased expression of toll-like receptor 2 on monocytes in HIV infection: possible roles in inflammation and viral replication. *Clin Infect Dis, 39*(2), 264-269. doi:10.1086/421780

Hernandez, J. C., Stevenson, M., Latz, E., & Urcuqui-Inchima, S. (2012). HIV type 1 infection up-regulates TLR2 and TLR4 expression and function in vivo and in vitro. *AIDS Res Hum Retroviruses, 28*(10), 1313-1328. doi:10.1089/aid.2011.0297

Hogerkorp, C. M., Nishimura, Y., Song, K., Martin, M. A., & Roederer, M. (2011). The simian immunodeficiency virus targets central cell cycle functions through transcriptional repression in vivo. *PLoS One, 6*(10), e25684. doi:10.1371/journal.pone.0025684

Homann, S., Smith, D., Little, S., Richman, D., & Guatelli, J. (2011). Upregulation of BST-2/Tetherin by HIV infection in vivo. *J Virol, 85*(20), 10659-10668. doi:10.1128/jvi.05524-11

Hotter, D., Bosso, M., Jonsson, K. L., Krapp, C., Sturzel, C. M., Das, A., . . . Kirchhoff, F. (2019). IFI16 Targets the Transcription Factor Sp1 to Suppress HIV-1 Transcription and Latency Reactivation. *Cell Host Microbe, 25*(6), 858-872.e813. doi:10.1016/j.chom.2019.05.002

Hubbard, J. J., Greenwell-Wild, T., Barrett, L., Yang, J., Lempicki, R. A., Wahl, S. M., . . . Kottilil, S. (2012). Host gene expression changes correlating with anti-HIV-1 effects in human subjects after treatment with peginterferon Alfa-2a. *J Infect Dis, 205*(9), 1443-1447. doi:10.1093/infdis/jis211

Irons, D. L., Meinhardt, T., Allers, C., Kuroda, M. J., & Kim, W. K. (2019). Overexpression and activation of colony-stimulating factor 1 receptor in the SIV/macaque model of HIV infection and neuroHIV. *Brain Pathol, 29*(6), 826-836. doi:10.1111/bpa.12731

Jaroenpool, J., Rogers, K. A., Pattanapanyasat, K., Villinger, F., Onlamoon, N., Crocker, P. R., & Ansari, A. A. (2007). Differences in the constitutive and SIV infection induced expression of Siglecs by hematopoietic cells from non-human primates. *Cell Immunol, 250*(1-2), 91-104. doi:10.1016/j.cellimm.2008.01.009

Kane, M., Yadav, S. S., Bitzegeio, J., Kutluay, S. B., Zang, T., Wilson, S. J., . . . Bieniasz, P. D. (2013). MX2 is an interferon-induced inhibitor of HIV-1 infection. *Nature, 502*(7472), 563-566. doi:10.1038/nature12653

Kim, S., Hahn, Y. K., Podhaizer, E. M., McLane, V. D., Zou, S., Hauser, K. F., & Knapp, P. E. (2018). A central role for glial CCR5 in directing the neuropathological interactions of HIV-1 Tat and opiates. *J Neuroinflammation, 15*(1), 285. doi:10.1186/s12974-018-1320-4

Knight, A. C., Brill, S. A., Queen, S. E., Tarwater, P. M., & Mankowski, J. L. (2018). Increased Microglial CSF1R Expression in the SIV/Macaque Model of HIV CNS Disease. *J Neuropathol Exp Neurol, 77*(3), 199-206. doi:10.1093/jnen/nlx115

Kunzi, M. S., & Pitha, P. M. (1996). Role of interferon-stimulated gene ISG-15 in the interferon-omega-mediated inhibition of human immunodeficiency virus replication. *J Interferon Cytokine Res, 16*(11), 919-927. doi:10.1089/jir.1996.16.919

Lederer, S., Favre, D., Walters, K. A., Proll, S., Kanwar, B., Kasakow, Z., . . . Katze, M. G. (2009). Transcriptional profiling in pathogenic and non-pathogenic SIV infections reveals significant distinctions in kinetics and tissue compartmentalization. *PLoS Pathog, 5*(2), e1000296. doi:10.1371/journal.ppat.1000296

Lenz, N., Schindler, T., Kagina, B. M., Zhang, J. D., Lukindo, T., Mpina, M., . . . Daubenberger, C. A. (2015). Antiviral Innate Immune Activation in HIV-Infected Adults Negatively Affects H1/IC31-Induced Vaccine-Specific Memory CD4+ T Cells. *Clin Vaccine Immunol, 22*(7), 688-696. doi:10.1128/cvi.00092-15

Li, Q., Smith, A. J., Schacker, T. W., Carlis, J. V., Duan, L., Reilly, C. S., & Haase, A. T. (2009). Microarray analysis of lymphatic tissue reveals stage-specific, gene expression signatures in HIV-1 infection. *J Immunol, 183*(3), 1975-1982. doi:10.4049/jimmunol.0803222

Li, W., Gorantla, S., Gendelman, H. E., & Poluektova, L. Y. (2017). Systemic HIV-1 infection produces a unique glial footprint in humanized mouse brains. *Dis Model Mech, 10*(12), 1489-1502. doi:10.1242/dmm.031773

Li, Y., Chan, E. Y., & Katze, M. G. (2007). Functional genomics analyses of differential macaque peripheral blood mononuclear cell infections by human immunodeficiency virus-1 and simian immunodeficiency virus. *Virology, 366*(1), 137-149. doi:10.1016/j.virol.2007.04.020

Lodermeyer, V., Ssebyatika, G., Passos, V., Ponnurangam, A., Malassa, A., Ewald, E., . . . Goffinet, C. (2018). The Antiviral Activity of the Cellular Glycoprotein LGALS3BP/90K Is Species Specific. *J Virol, 92*(14). doi:10.1128/jvi.00226-18

Lorenz, D. R., Misra, V., & Gabuzda, D. (2019). Transcriptomic analysis of monocytes from HIV-positive men on antiretroviral therapy reveals effects of tobacco smoking on interferon and stress response systems associated with depressive symptoms. *Hum Genomics, 13*(1), 59. doi:10.1186/s40246-019-0247-x

Lu, W., Demers, A. J., Ma, F., Kang, G., Yuan, Z., Wan, Y., . . . Li, Q. (2016). Next-Generation mRNA Sequencing Reveals Pyroptosis-Induced CD4+ T Cell Death in Early Simian Immunodeficiency Virus-Infected Lymphoid Tissues. *J Virol, 90*(2), 1080-1087. doi:10.1128/jvi.02297-15

Lu, W., Ma, F., Churbanov, A., Wan, Y., Li, Y., Kang, G., . . . Li, Q. (2014). Virus-host mucosal interactions during early SIV rectal transmission. *Virology, 464-465*, 406-414. doi:10.1016/j.virol.2014.07.010

Maingat, F., Halloran, B., Acharjee, S., van Marle, G., Church, D., Gill, M. J., . . . Power, C. (2011). Inflammation and epithelial cell injury in AIDS enteropathy: involvement of endoplasmic reticulum stress. *Faseb j, 25*(7), 2211-2220. doi:10.1096/fj.10-175992

Matheson, N. J., Sumner, J., Wals, K., Rapiteanu, R., Weekes, M. P., Vigan, R., . . . Lehner, P. J. (2015). Cell Surface Proteomic Map of HIV Infection Reveals Antagonism of Amino Acid Metabolism by Vpu and Nef. *Cell Host Microbe, 18*(4), 409-423. doi:10.1016/j.chom.2015.09.003

Mohan, M., Kaushal, D., Aye, P. P., Alvarez, X., Veazey, R. S., & Lackner, A. A. (2013). Focused examination of the intestinal epithelium reveals transcriptional signatures consistent with disturbances in enterocyte maturation and differentiation during the course of SIV infection. *PLoS One, 8*(4), e60122. doi:10.1371/journal.pone.0060122

Mothapo, K. M., Ten Oever, J., Koopmans, P., Stelma, F. F., Burm, S., Bajramovic, J., . . . van der Ven, A. J. (2017). Soluble TLR2 and 4 concentrations in cerebrospinal fluid in HIV/SIV-related neuropathological conditions. *J Neurovirol, 23*(2), 250-259. doi:10.1007/s13365-016-0495-7

Mussil, B., Javed, A., Topfer, K., Sauermann, U., & Sopper, S. (2015). Increased BST2 expression during simian immunodeficiency virus infection is not a determinant of disease progression in rhesus monkeys. *Retrovirology, 12*, 92. doi:10.1186/s12977-015-0219-8

Nasr, N., Alshehri, A. A., Wright, T. K., Shahid, M., Heiner, B. M., Harman, A. N., . . . Cunningham, A. L. (2017). Mechanism of Interferon-Stimulated Gene Induction in HIV-1-Infected Macrophages. *J Virol, 91*(20). doi:10.1128/jvi.00744-17

Nowlin, B. T., Wang, J., Schafer, J. L., Autissier, P., Burdo, T. H., & Williams, K. C. (2018). Monocyte subsets exhibit transcriptional plasticity and a shared response to interferon in SIV-infected rhesus macaques. *J Leukoc Biol, 103*(1), 141-155. doi:10.1002/jlb.4a0217-047r

Omi, K., Shimizu, M., Watanabe, E., Matsumura, J., Takaku, C., Shinya, E., & Takahashi, H. (2014). Inhibition of R5-tropic HIV type-1 replication in CD4(+) natural killer T cells by gammadelta T lymphocytes. *Immunology, 141*(4), 596-608. doi:10.1111/imm.12221

Pallikkuth, S., Parmigiani, A., & Pahwa, S. (2012). Role of IL-21 and IL-21 receptor on B cells in HIV infection. *Crit Rev Immunol, 32*(2), 173-195. doi:10.1615/critrevimmunol.v32.i2.50

Paparisto, E., Woods, M. W., Coleman, M. D., Moghadasi, S. A., Kochar, D. S., Tom, S. K., . . . Barr, S. D. (2018). Evolution-Guided Structural and Functional Analyses of the HERC Family Reveal an Ancient Marine Origin and Determinants of Antiviral Activity. *J Virol, 92*(13). doi:10.1128/jvi.00528-18

Periyasamy, P., Thangaraj, A., Bendi, V. S., & Buch, S. (2019). HIV-1 Tat-mediated microglial inflammation involves a novel miRNA-34a-NLRC5-NFkappaB signaling axis. *Brain Behav Immun, 80*, 227-237. doi:10.1016/j.bbi.2019.03.011

Rempel, H., Sun, B., Calosing, C., Abadjian, L., Monto, A., & Pulliam, L. (2013). Monocyte activation in HIV/HCV coinfection correlates with cognitive impairment. *PLoS One, 8*(2), e55776. doi:10.1371/journal.pone.0055776

Roberts, E. S., Zandonatti, M. A., Watry, D. D., Madden, L. J., Henriksen, S. J., Taffe, M. A., & Fox, H. S. (2003). Induction of pathogenic sets of genes in macrophages and neurons in NeuroAIDS. *Am J Pathol, 162*(6), 2041-2057. doi:10.1016/s0002-9440(10)64336-2

Ross, C. T., Roodgar, M., & Smith, D. G. (2015). Evolutionary distance of amino acid sequence orthologs across macaque subspecies: identifying candidate genes for SIV resistance in Chinese rhesus macaques. *PLoS One, 10*(4), e0123624. doi:10.1371/journal.pone.0123624

Rotger, M., Dalmau, J., Rauch, A., McLaren, P., Bosinger, S. E., Martinez, R., . . . Telenti, A. (2011). Comparative transcriptomics of extreme phenotypes of human HIV-1 infection and SIV infection in sooty mangabey and rhesus macaque. *J Clin Invest, 121*(6), 2391-2400. doi:10.1172/jci45235

Sabado, R. L., O'Brien, M., Subedi, A., Qin, L., Hu, N., Taylor, E., . . . Bhardwaj, N. (2010). Evidence of dysregulation of dendritic cells in primary HIV infection. *Blood, 116*(19), 3839-3852. doi:10.1182/blood-2010-03-273763

Sandler, N. G., Bosinger, S. E., Estes, J. D., Zhu, R. T., Tharp, G. K., Boritz, E., . . . Douek, D. C. (2014). Type I interferon responses in rhesus macaques prevent SIV infection and slow disease progression. *Nature, 511*(7511), 601-605. doi:10.1038/nature13554

Sanfilippo, C., Pinzone, M. R., Cambria, D., Longo, A., Palumbo, M., Di Marco, R., . . . Di Rosa, M. (2018). OAS Gene Family Expression Is Associated with HIV-Related Neurocognitive Disorders. *Mol Neurobiol, 55*(3), 1905-1914. doi:10.1007/s12035-017-0460-3

Shiau, S., Strehlau, R., Wang, S., Violari, A., Do, C., Patel, F., . . . Kuhn, L. (2019). Distinct epigenetic profiles in children with perinatally-acquired HIV on antiretroviral therapy. *Sci Rep, 9*(1), 10495. doi:10.1038/s41598-019-46930-1

Siangphoe, U., & Archer, K. J. (2015). Gene Expression in HIV-Associated Neurocognitive Disorders: A Meta-Analysis. *J Acquir Immune Defic Syndr, 70*(5), 479-488. doi:10.1097/qai.0000000000000800

Stewart, J. C., Polanka, B. M., So-Armah, K. A., White, J. R., Gupta, S. K., Kundu, S., . . . Freiberg, M. S. (2020). Associations of Total, Cognitive/Affective, and Somatic Depressive Symptoms and Antidepressant Use with Cardiovascular Disease-Relevant Biomarkers in HIV: Veterans Aging Cohort Study. *Psychosom Med*. doi:10.1097/psy.0000000000000808

Tremblay-McLean, A., Bruneau, J., Lebouche, B., Lisovsky, I., Song, R., & Bernard, N. F. (2017). Expression Profiles of Ligands for Activating Natural Killer Cell Receptors on HIV Infected and Uninfected CD4(+) T Cells. *Viruses, 9*(10). doi:10.3390/v9100295

Trifone, C., Salido, J., Ruiz, M. J., Leng, L., Quiroga, M. F., Salomon, H., . . . Turk, G. (2018). Interaction Between Macrophage Migration Inhibitory Factor and CD74 in Human Immunodeficiency Virus Type I Infected Primary Monocyte-Derived Macrophages Triggers the Production of Proinflammatory Mediators and Enhances Infection of Unactivated CD4(+) T Cells. *Front Immunol, 9*, 1494. doi:10.3389/fimmu.2018.01494

Turrini, F., Saliu, F., Forlani, G., Das, A. T., Van Lint, C., Accolla, R. S., . . . Vicenzi, E. (2019). Interferon-inducible TRIM22 contributes to maintenance of HIV-1 proviral latency in T cell lines. *Virus Res, 269*, 197631. doi:10.1016/j.virusres.2019.05.009

Vitalle, J., Tarancon-Diez, L., Jimenez-Leon, M. R., Terren, I., Orrantia, A., Roca-Oporto, C., . . . Borrego, F. (2020). CD300a identifies a CD4+ memory T cell subset with a higher susceptibility to HIV-1 infection. *Aids*. doi:10.1097/qad.0000000000002544

Vlasova-St Louis, I., Chang, C. C., Shahid, S., French, M. A., & Bohjanen, P. R. (2018). Transcriptomic Predictors of Paradoxical Cryptococcosis-Associated Immune Reconstitution Inflammatory Syndrome. *Open Forum Infect Dis, 5*(7), ofy157. doi:10.1093/ofid/ofy157

Wachter, C., Eiden, L. E., Naumann, N., Depboylu, C., & Weihe, E. (2016). Loss of cerebellar neurons in the progression of lentiviral disease: effects of CNS-permeant antiretroviral therapy. *J Neuroinflammation, 13*(1), 272. doi:10.1186/s12974-016-0726-0

Wang, L., Huang, J., Jiang, M., & Sun, L. (2011). MYBPC1 computational phosphoprotein network construction and analysis between frontal cortex of HIV encephalitis (HIVE) and HIVE-control patients. *Cell Mol Neurobiol, 31*(2), 233-241. doi:10.1007/s10571-010-9613-x

Wie, S. H., Du, P., Luong, T. Q., Rought, S. E., Beliakova-Bethell, N., Lozach, J., . . . Woelk, C. H. (2013). HIV downregulates interferon-stimulated genes in primary macrophages. *J Interferon Cytokine Res, 33*(2), 90-95. doi:10.1089/jir.2012.0052

Williams, M. E., Ipser, J. C., Stein, D. J., Joska, J. A., & Naude, P. J. W. (2019). The Association of Immune Markers with Cognitive Performance in South African HIV-Positive Patients. *J Neuroimmune Pharmacol, 14*(4), 679-687. doi:10.1007/s11481-019-09870-1

Winkler, J. M., Chaudhuri, A. D., & Fox, H. S. (2012). Translating the brain transcriptome in neuroAIDS: from non-human primates to humans. *J Neuroimmune Pharmacol, 7*(2), 372-379. doi:10.1007/s11481-012-9344-5

Woods, M. W., Kelly, J. N., Hattlmann, C. J., Tong, J. G., Xu, L. S., Coleman, M. D., . . . Barr, S. D. (2011). Human HERC5 restricts an early stage of HIV-1 assembly by a mechanism correlating with the ISGylation of Gag. *Retrovirology, 8*, 95. doi:10.1186/1742-4690-8-95

Wu, J. Q., Sasse, T. R., Saksena, M. M., & Saksena, N. K. (2013). Transcriptome analysis of primary monocytes from HIV-positive patients with differential responses to antiretroviral therapy. *Virol J, 10*, 361. doi:10.1186/1743-422x-10-361

Yang, Z., Yang, J., Wang, J., Lu, X., Jin, C., Xie, T., & Wu, N. (2015). Identify Potential Regulators in HIV-1 Latency by Joint microRNA and mRNA Analysis. *Cell Physiol Biochem, 36*(2), 569-584. doi:10.1159/000430121

Zahoor, M. A., Xue, G., Sato, H., & Aida, Y. (2015). Genome-wide transcriptional profiling reveals that HIV-1 Vpr differentially regulates interferon-stimulated genes in human monocyte-derived dendritic cells. *Virus Res, 208*, 156-163. doi:10.1016/j.virusres.2015.06.017

Zahoor, M. A., Xue, G., Sato, H., Murakami, T., Takeshima, S. N., & Aida, Y. (2014). HIV-1 Vpr induces interferon-stimulated genes in human monocyte-derived macrophages. *PLoS One, 9*(8), e106418. doi:10.1371/journal.pone.0106418

Zhang, W., Ambikan, A. T., Sperk, M., van Domselaar, R., Nowak, P., Noyan, K., . . . Neogi, U. (2018). Transcriptomics and Targeted Proteomics Analysis to Gain Insights Into the Immune-control Mechanisms of HIV-1 Infected Elite Controllers. *EBioMedicine, 27*, 40-50. doi:10.1016/j.ebiom.2017.11.031

Zhou, M., Greenhill, S., Huang, S., Silva, T. K., Sano, Y., Wu, S., . . . Silva, A. J. (2016). CCR5 is a suppressor for cortical plasticity and hippocampal learning and memory. *Elife, 5*. doi:10.7554/eLife.20985

Zhou, Y., Rong, L., Lu, J., Pan, Q., & Liang, C. (2008). Insulin-like growth factor II mRNA binding protein 1 associates with Gag protein of human immunodeficiency virus type 1, and its overexpression affects virus assembly. *J Virol, 82*(12), 5683-5692. doi:10.1128/jvi.00189-08
